# Context-dependent gene regulation by transcription factor complexes

**DOI:** 10.1101/706473

**Authors:** Judith F. Kribelbauer, Ryan E. Loker, Siqian Feng, Chaitanya Rastogi, Namiko Abe, H. Tomas Rube, Harmen J. Bussemaker, Richard S. Mann

## Abstract

Eukaryotic transcription factors (TFs) form complexes with various partner proteins to recognize their genomic target sites. Yet, how the DNA sequence determines which TF complex forms at any given site is poorly understood. Here we demonstrate that high-throughput *in vitro* binding assays coupled with unbiased computational analysis provides unprecedented insight into how complexes of homeodomain proteins adapt their stoichiometry and configuration to the bound DNA. Using inferred knowledge about minor groove width readout, we design targeted protein mutations that destabilize homeodomain binding in a complex-specific manner. By performing parallel SELEX-seq, ChIP-seq, RNA-seq and Hi-C assays, we not only reveal complex-specific functions, but also show that TF binding sites that lack a canonical sequence motif emerge as a consequence of direct interaction with functionally bound sites.

## INTRODUCTION

Gene regulatory networks are controlled by transcription factors (TFs) that target distinct gene sets by binding to specific DNA sequences. To determine which genes are regulated by a given TF, the genome-wide pattern of TF binding must be assayed and interpreted. The current standard approach is to profile *in vivo* TF occupancy using ChIP-seq or related methods(*1*–*5*). However, because these assays are blind to which co-factors the TF uses to bind any particular locus, it is difficult to infer how the DNA sequence determines the stoichiometry and composition of TF complex assembly.

A complementary approach to identify TF binding sites involves probing the DNA binding specificity of TFs using high-throughput *in vitro* assays (*6*). Binding preferences derived from such experiments are typically summarized by a position weight matrix or PWM (*7*). Despite their popularity, PWMs typically fail to explain a large fraction of *in vivo* TF binding events in higher eukaryotes (*8*). There are several possible explanations for this: For one, low-affinity binding sites, which generally do not harbor a motif match, can be bound and functional *in vivo* (*9*, *10*). Second, a TF may bind its genomic target sites cooperatively with other TFs (*11*–*15*) or with nucleosomes (*16*). Finally, indirect pull-down at highly accessible sites (*17*) or experimental artifacts (*18*) may also contribute. Although the mechanism by which ChIP enrichment is accrued in the absence of sequence-specific binding is not well understood, recent insights into the compartmentalized structure of eukaryotic nuclei and the formation of transcriptional hubs with high concentration of TFs (*19*, *20*) provide a potential explanation.

Yet another approach to analyzing TF binding specificity is to obtain atomic-resolution structural information of protein-DNA complexes. To date, the structures of several thousands of protein-nucleic acid complexes have been determined (*21*), including representatives for all major TF families (*22*). However, as with PWM models, the majority of these structures were obtained using only the DNA binding domain (DBD) bound to a single DNA ligand and, as a result, provide little structural insight into the range of binding modes exhibited by combinations of full-length TFs *in vivo*.

Despite their individual short-comings, these different approaches have yielded a rich trove of complementary data sets, which together may allow us to dissect the causal relationship between DNA sequence, TF complex binding, nuclear transcriptional hub assembly, and coordinated regulation of gene expression.

Here we combine high-throughput *in vitro* binding experiments with structural information to design engineered TFs that can elucidate important principles underlying gene regulatory networks. Importantly, by contrasting the molecular behavior of “wild-type” and “engineered” versions of the same TF *in vivo*, we obtain detailed information on TF complex-specific gene control and function. To illustrate this approach, we focused on a system of three interacting homeodomain (HD) transcription factors from *D. melanogaster* – one of the eight Hox proteins in the presence of the homeodomain cofactors Homothorax (Hth) and Extradenticle (Exd). This TF system exhibits many of the complexities that exist for most eukaryotic TFs, including overlapping binding specificities within large TF families (*23*, *24*), the existence of multiple TF isoforms (*25*, *26*), cooperativity and latent DNA binding specificities (*13*), and distinct biological functions that depend on unique TF complex compositions (*25*, *27*–*29*).

We show that, as with Hox homeodomains (*30*), basic amino acids within the N-terminal arms of both Hth and Exd homeodomains prefer to bind DNA sequences with narrow minor grooves. We use this insight to design targeted protein mutations that are compromised in this mode of recognition and, as a consequence, selectively eliminate the binding of some, but not all, Exd-containing complexes. Exploiting this differential sensitivity as a tool *in vivo*, we classify each binding site according to the specific homeodomain complex that it binds. Furthermore, by combining information on 3D chromatin interactions with the variable degree of mutant Exd binding loss, we demonstrate that binding to sites lacking a sequence motif results from direct interactions with sites bound in a sequence-specific manner. Finally, we infer hidden complex-specific biological functions by linking distinct Exd complexes to the set of target promoters they physically interact with.

## RESULTS

### Hox, Hth, and Exd form complexes with distinct sequence and conformation preferences

The TALE-family homeobox protein Exd can form a heterodimer with each of the eight *D. melanogaster* Hox factors (*13*). Nuclear localization of Exd is dependent on Hth (*31*, *32*), a second TALE-family homeobox TF that exists as two major isoforms (*25*): a full-length, homeodomain (HD) containing isoform, Hth^FL^, and a shorter, HD-less isoform, Hth^HM^ (Homothorax-Meis domain). Since the tight Exd-Hth protein-protein interaction occurs between the HM domain and Exd’s PBC domain (*33*) (**Fig. 1A**), both isoforms are sufficient for the nuclear localization of Exd. In addition to acting as a Hox cofactor, Hth^FL^-Exd carries out Hox-independent functions such as patterning the proximal-distal axes of the appendages and specifying antennal identity (*34*–*36*). As a result, a variety of Exd-containing complexes are present *in vivo* – Hth^FL^-Exd-Hox, with three HDs, Hth^HM^-Exd-Hox or Hth^FL^-Exd, each with two HDs – as well as Hth^FL^ binding as a monomer or homodimer without direct Exd-DNA contact (**Fig. 1B**). Structural information, however, is largely limited to the HDs of heterodimeric Exd-Hox and homodimeric MEIS1 (the human ortholog of Hth) (**Fig. 1B**). Thus, it remains unclear how the assembly of different complexes is promoted by the DNA sequence, or how the combinatorial nature of homeodomain binding contributes to gene regulation.

**Fig. 1:**
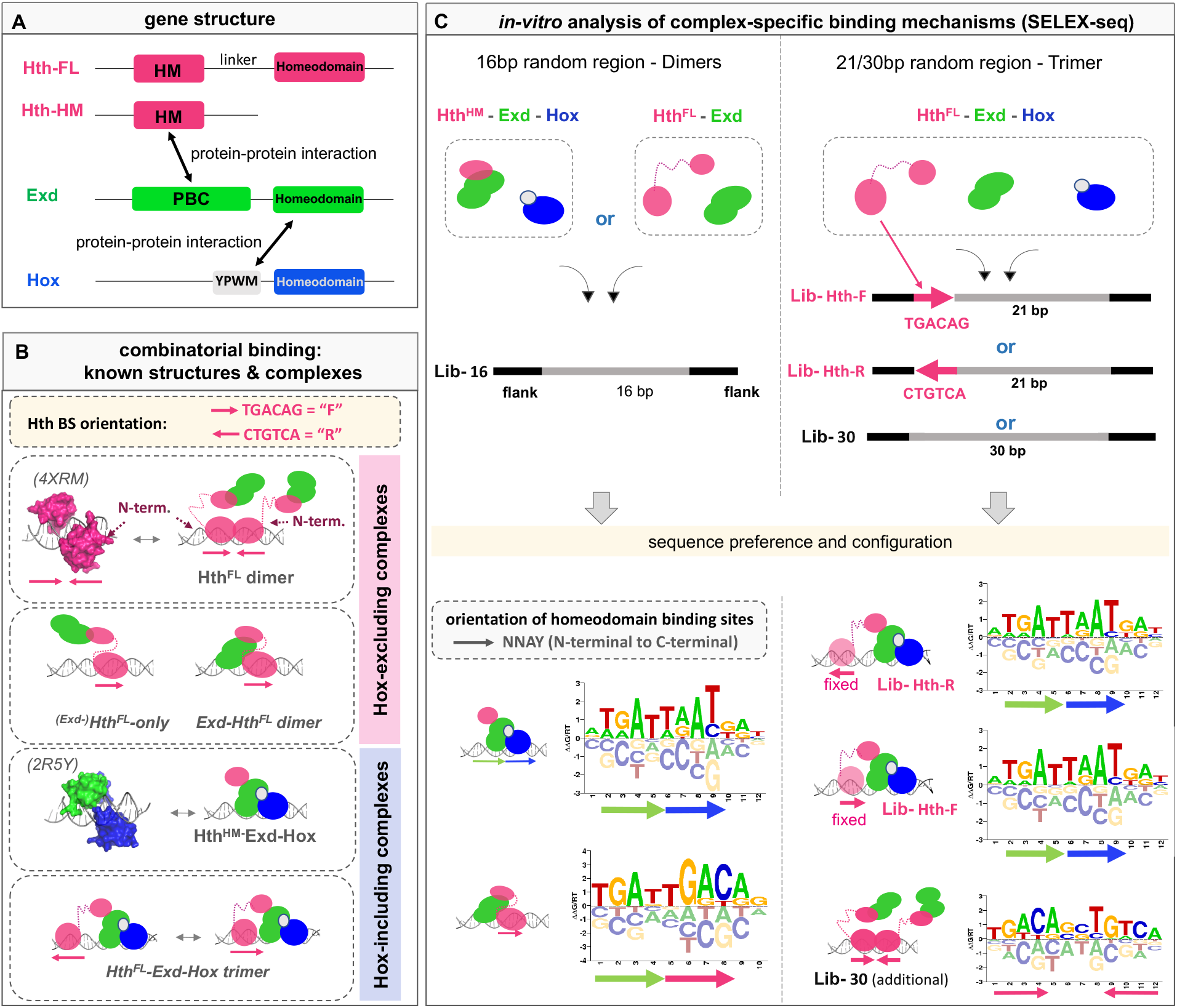
Probing the diversity of multi-homeodomain complex binding using SELEX-seq. **(A)** Schematic gene structures for three homeodomain TFs: Homothorax (Hth; pink), Extradenticle (Exd; green), Hox (Hox; blue). Arrows indicate protein interactions: PBC-domain (PBC); Homothorax-Meis domain (HM); YPWM (Exd interaction motif). **(B)** Existing 3D structures and schematic diagrams showing various possible complexes formed by Hth, Exd, and/or Hox. Arrows indicate Hth binding site (BS) orientation (forward, F = TGACAG and reverse, R = CTGTCA). **(C)** SELEX-seq library design and derived sequence motifs (shown as energy logos). Arrows indicate protein binding site orientation with respect to the consensus NNAY homeodomain motif.

To characterize *in vitro* binding preferences, we designed SELEX-seq libraries (*13*, *37*, *38*) whose randomized region can accommodate the entire footprint of each respective complex (**Fig. 1C**). To facilitate analysis of complex binding patterns when all three HDs are present, we designed two libraries in which a fixed Hth binding site immediately precedes a 21-bp randomized region: (i) Lib-Hth-F, with an Hth site (TGACAG) designed to bind Hth in <u>f</u>orward orientation, and (ii) Lib-Hth-R, with a <u>r</u>everse site (CTGTCA) (**Fig. 1C**). We carried out SELEX-seq experiments for all individual complexes and constructed position-specific affinity matrices (PSAMs) and energy logos (*39*) based on the relative enrichment of oligomers of a given length (**Fig. 1C**; see **Methods**). This analysis indicates that in the absence of Hox, Exd-Hth^FL^ prefers to bind as a head-to-tail dimer analogous to Exd-Hox (**Fig. 1C**). Introducing a Hox protein to the Hth^FL^-Exd complex results in the formation of a dominant Exd-Hox subcomplex, similar to when orientation-agnostic libraries are used (**Fig. S1A,B**). Sequences suggestive of Exd-Hth^FL^ (dark blue) and Hth^FL^–Hth^FL^ dimer binding (dark pink) are also observed (**Fig. S1B**).

### Relative position and orientation preferences of a ternary protein-DNA complex

Characterizing the binding preferences of the ternary Hth^FL^-Exd-Hox complex requires taking into consideration both the orientation and position of the Hth^FL^ binding site relative to the Exd-Hox heterodimer binding site. To infer this information from the SELEX-seq data, we first computed the relative enrichment of all DNA 12-mers both for Hth^FL^-Exd-Hox and Hth^HM^-Exd-Hox (Fig. **S1A,B**). Using the PSAM for Exd-Hox to assign binding orientation, we find that in the absence of a Hth homeodomain, similar enrichments are observed for the forward ([Exd-Hox]_F_) and reverse ([Exd-Hox]_R_) orientations. However, when the homeodomain-containing isoform Hth^FL^ is used, the configuration [Hth^FL^]_F_[Exd-Hox]_F_ is preferred over [Hth^FL^]_F_[Exd-Hox]_R_ (Fig. **S1A, B & C**).

Next, we estimated the contribution to the total binding free energy associated with the “full configuration” (i.e., the relative position and orientation of the Hth and Exd-Hox subunits) by fitting a generalized linear model (GLM; see **Methods**) (**Fig. 2A**). For both Hth binding site orientations (F and R), the configuration in which Hth binds on the Exd side of the Exd-Hox dimer was favored. In addition, a preference for shorter spacers was observed for the Hth-F library compared to the Hth-R library (**Fig. 2A** and **Fig. S1D**). This preference suggests that the N-terminus of Hth’s HD faces Exd in Lib-Hth-R, shortening the distance between Hth’s HM and Exd’s PBC domains and thus allowing for a longer DNA spacer, while facing away in Lib-Hth-F, requiring the Exd-Hox subcomplex to move closer to the Hth binding site. The proposed structural configuration indeed makes mechanistic sense in light of the MEIS1 crystal structure (the human Hth ortholog; PDB-ID: 4XRM) (**Fig. 2B**) (*15*).

**Fig. 2:**
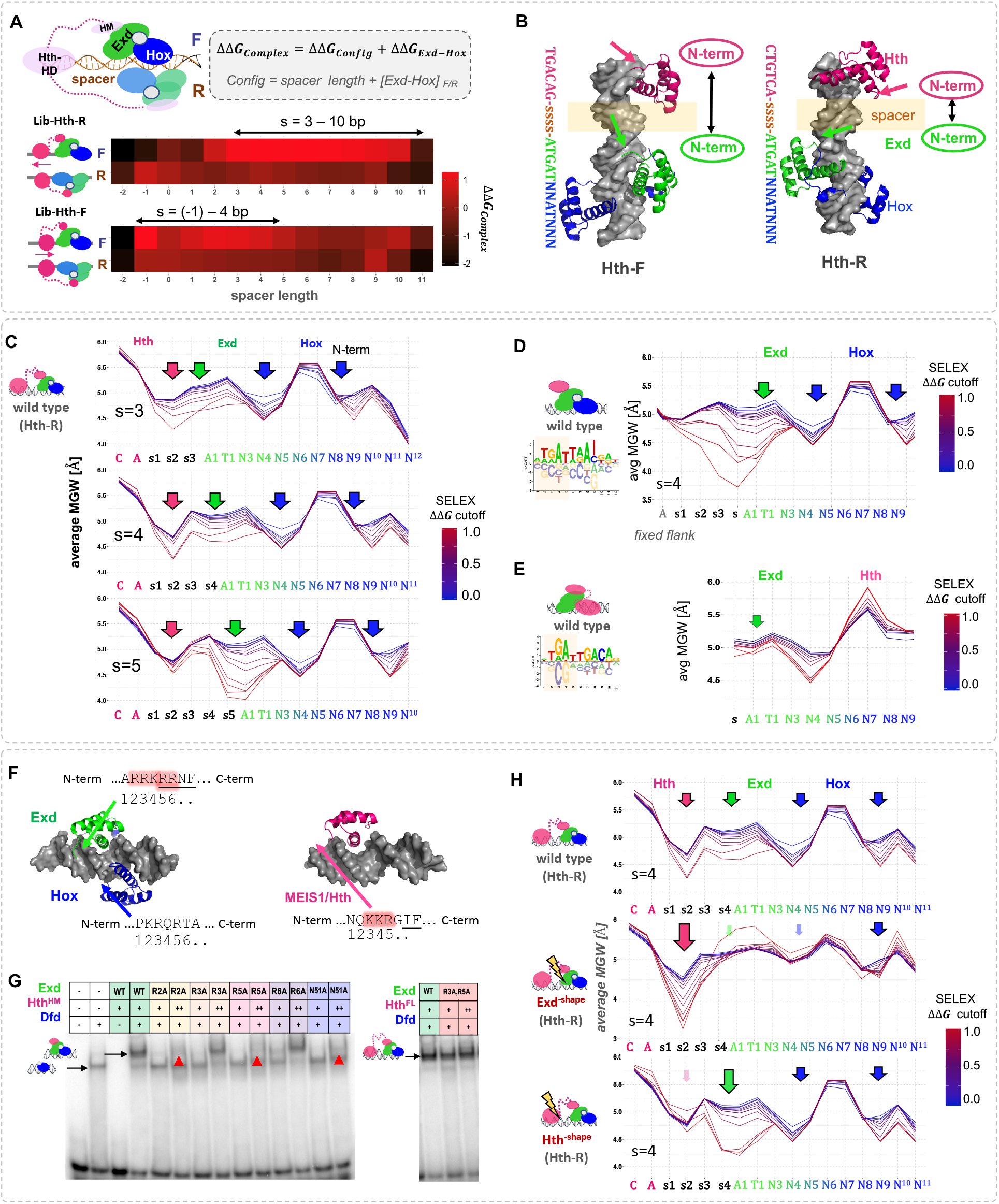
Dissecting DNA minor groove width readout by a ternary homeodomain complex. **(A)** Systematic analysis of binding configurations of the ternary Hth^FL^-Exd-Hox complex. SELEX probe counts after two rounds of affinity-based selection were analyzed using a generalized linear model that estimates the free energy associated with each configuration (i.e., the length of DNA spacer between the Hth and Exd-Hox sites, and their orientation with respect to each other) while accounting for the dependence on DNA sequence based on the enrichment of 12-mers observed for the simpler Hth^HM^-Exd-Hox complex. Heatmaps show ΔΔ*G* coefficients (in units of RT) for each particular configuration; red indicates stronger binding. (B) Superposition of Meis1 (human ortholog of Hth; PDB-ID: 4XRM) and Exd-Hox (PDB-ID: 2R5Y) crystal structures onto B-DNA templates (http://structure.usc.edu/make-na/) consisting of a Hth-F (TGACAG) or Hth-R (CTGTCA) binding site, followed by a 4-bp spacer (indicated by “ssss”) and an Exd-Hox site (2R5Y). Arrows indicate the relative positioning of the N-terminal domain of each HD (Hth: pink; Exd: green). **(C-E)** Profiles of average minor groove width (MGW) at increasingly stringent affinity cutoffs, for **(C)** Hth^FL^-Exd-Hox (Lib-Hth-R) and three different spacer lengths (3-5 bp), **(D)** Hth^HM^-Exd-Hox, with spacer of 4 bp, and **(E)** Hth^FL^-Exd. Arrows indicate MGW minima. **(F)** Crystal structures of Exd-Hox heterodimer (PDB-ID:2R5Y) and Meis1 homodimer (PDB-ID:4XRM; human ortholog of Hth) with the sequence of their N-terminal HD arms indicated; red shading indicates positively charged amino acids that were mutated in this study. Amino acids are partially (Exd) or entirely (Hth) unresolved in the crystal structures (resolved amino acids are underlined). **(G)** Electromobility shift assay (EMSA) for single and double amino-acid point mutations of Exd-Hox (with Dfd playing the role of Hox) in complex with either Hth^HM^ or Hth^FL^. Arrows indicate binding loss. **(H)** Verification of MGW readout using shape-defective mutants of Hth (middle row) or Exd (bottom row). Shaded arrows indicate the loss of specific MGW minima.

To validate our configurational free energy estimates, we performed competition electromobility shift assays (EMSAs) on three different DNA spacer lengths (**Fig. S1E**).

### Homeodomain complexes vary in their dependency on the recognition of DNA shape

The currently available structures of homeodomain-DNA complexes suggest that the spacer DNA separating the Hth^FL^ and Exd-Hox binding sites is not directly contacted by any of these proteins. However, since DBDs were used rather than full-length proteins, the contacts observed in these structures may not capture all relevant contributions to complex stability. To determine whether the sequence of the DNA spacer might contribute to the thermodynamic stability of the complex, we computed oligomer enrichment over the first four nucleotide positions downstream of the fixed Hth site in Lib-Hth-R, retaining only those probes that matched the 12-bp PSAM for Exd-Hox over positions 5-16. A preference for AT-rich sequences observed in the most highly enriched spacers (**Fig. S1F**) suggested that the spacer may influence binding affinity via DNA minor groove width (MGW) readout, which has been shown to play a critical role in DNA recognition for many TFs (*15*, *30*, *40*–*42*).

To analyze the relationship between spacer sequence preference and DNA shape readout, we fit a mechanism-agnostic GLM based on base identities over the first 15 nucleotide positions of the variable region (3-bp spacer and a 12-bp Exd-Hox site), keeping the first two base pairs within the Exd-Hox site fixed (**Fig. S1G,H**; see **Methods** for details). Consistent with recent analyses (Rube et al., 2018), spacer preferences derived from a GLM that neglects dependencies between nucleotide positions agreed well with a GLM in which each spacer oligonucleotide was scored separately (R^2^=0.81, **Fig. S1I**). By taking subsets of sequences defined by an increasingly stringent cutoff on their predicted overall affinity and computing their average minor groove width (MGW) using pentamer tables (*43*), we visualized the relationship between intrinsic DNA shape and probe selection in the SELEX-seq assay (**Fig. 2C and Fig. S1G**). In addition to the two known MGW minima preferred by anterior Hox TFs (*30*, *44*), we observe a strong preference for a narrow minor groove within the 3-bp spacer region separating the Hth and Exd-Hox binding sites. Notably, when we used the same approach to examine spacers longer than 3 bp, this broad MGW minimum split into two narrow ones anchored to the Hth and Exd binding sites, respectively (**Fig. 2C**).

These observations suggest that both Hth and Exd can take advantage of local narrowing of the DNA minor groove by directing positively charged amino acids within their N-terminal arms towards the DNA spacer (**Fig. 2C**). To test this hypothesis, we repeated the analysis for Hth^HM^-Exd-Hox and Exd-Hth^FL^, reasoning that the Exd MGW minimum would be observed for all three complexes, whereas the Hth-associated one would be absent in the Hth^HM^ complex. Indeed, we detected the Exd-associated, but not the Hth^FL^-associated MGW minimum for the Hth^HM^-Exd-Hox complex (**Fig. 2D**). Contrary to expectation, Exd’s strong preference for a narrow minor groove is attenuated in the Exd-Hth^FL^ complex (**Fig. 2E**), despite that fact that the same Exd site (ATGAT) is optimal for both complexes. Intriguingly, these observations suggest that by analyzing the sequence selection trends in a SELEX experiment in terms of DNA shape features, it is possible to infer structural readout mechanisms specific to a particular TF complex that would otherwise remain elusive.

### Engineered mutations disrupting shape readout lead to selective loss of complex binding

To verify that the positively-charged residues in the N-terminal arms of Hth and Exd play a role in complex-specific DNA shape readout, we created mutants with up to three arginine/lysine (R/K) to alanine (A) substitutions in the N-terminal arms of either the Exd or Hth^FL^ HD, and performed EMSAs for Hth^HM^-Exd-Hox, Hth^FL^-Exd-Hox and Exd-Hth^FL^ complexes (**Fig. 2F,G** and **Fig. S2**). Strikingly, two single substitutions within Exd’s N-terminal arm (Exd^R2A^ and Exd^R5A^) were each sufficient to abrogate binding of the Hth^HM^-Exd-Hox complex to the same extent as a key hydrogen bonded residue in the α3 recognition helix of Exd (Exd^N51A^; for numbering of amino acids see (*24*)). When the same R-to-A mutations in Exd were tested in the context of Hth^FL^-Exd-Hox or Exd-Hth^FL^, binding of the complex was only mildly affected (**Fig. 2G** and **Fig. S2**). Similarly, a triple mutant Hth^K3A,K4A,R5A^ was still capable of binding DNA when complexed with Exd-Hox. However, weaker trimer binding and stronger monomer binding was observed when both N-terminal arms were mutated (Hth^K3A,K4A,R5A^-Exd^R3A,R5A^-Dfd) (**Fig. S2**). These findings demonstrate that although these contacts are not visible in existing crystal structures (**Fig. 2F**), the N-terminal arms of both Exd and Hth contribute extensively to binding. Importantly, their requirement is context dependent: the three-HD Hth^FL^-Exd-Hox and the two-HD Exd-Hth^FL^ complex can tolerate mutations in either the Hth or Exd N-terminal arm, while the two-HD Hth^HM^-Exd-Hox complex cannot.

To assess the impact of mutating these N-terminal arm residues across all binding sites, we performed SELEX-seq assays with mutant Hth^FL^-Exd-Hox and Exd-Hth^FL^ complexes, containing either Hth^K3A,K4A,R5A^ (which we will refer to as Hth^−shape^) or Exd^R2A,R5A^ (which we refer to as Exd^−shape^). Repeating the analysis of Fig. 2C revealed that the mutant Hth^FL^-Exd-Hox complexes lost the ability to select sequences containing the corresponding MGW minima (**Fig. 2H**). In addition, Exd^−shape^ blunted one of the previously observed (*13*, *30*, *44*) preferences for a narrow minor groove within the Exd-Hox binding site, suggesting that Exd-driven shape-readout promotes Exd-Hox heterodimer binding. Strikingly, this effect was not observed for Exd^−shape^-Hth^FL^ heterodimers; when compared to the wild-type Exd-Hth^FL^ complex, only the N-terminal Exd MGW readout was mildly perturbed, yet the MGW sensitivity at the first AY (position 4/5) remained (**Fig. S3A**). Therefore, the mutant-Exd data confirms our prediction that Exd’s N-terminal arm engages with the minor groove to a different degree when bound with Hth instead of Hox.

### Differential sensitivity to shape readout mutation discriminates between TF complexes

To determine the extent to which the Exd shape-readout mutation differentially impacts complex formation, we systematically compared the sequences selected by Hth^FL^-Exd-Hox with those selected by Hth^FL^-Exd^−shape^-Hox complexes (**Fig. 3A** and **Fig. S4D**). This analysis revealed large variation in the extent to which binding to a particular DNA sequence is affected by the loss of DNA shape readout by Exd. In an attempt to interpret these observations, we used the PSAMs derived for each distinct complex (cf. **Fig. 1C**) to predict the identity of the bound complex for each DNA sequence (colors in **Fig. 3A** and **Fig. S4D, G**). This analysis confirmed that in addition to discriminating between Hth-Exd-Hox complexes containing the Hth^FL^ and Hth^HM^ isoform respectively, Exd^−shape^ separates Hox-containing from non-Hox-containing complexes (**Fig. 3A** and **Fig. S4D, G**). Binding of Hth^FL^ homodimers was not impacted, binding of Exd-Hth^FL^ heterodimers was slightly impacted, and binding of Hth^FL^-Exd-Hox ternary complexes was strongly affected.

**Fig. 3:**
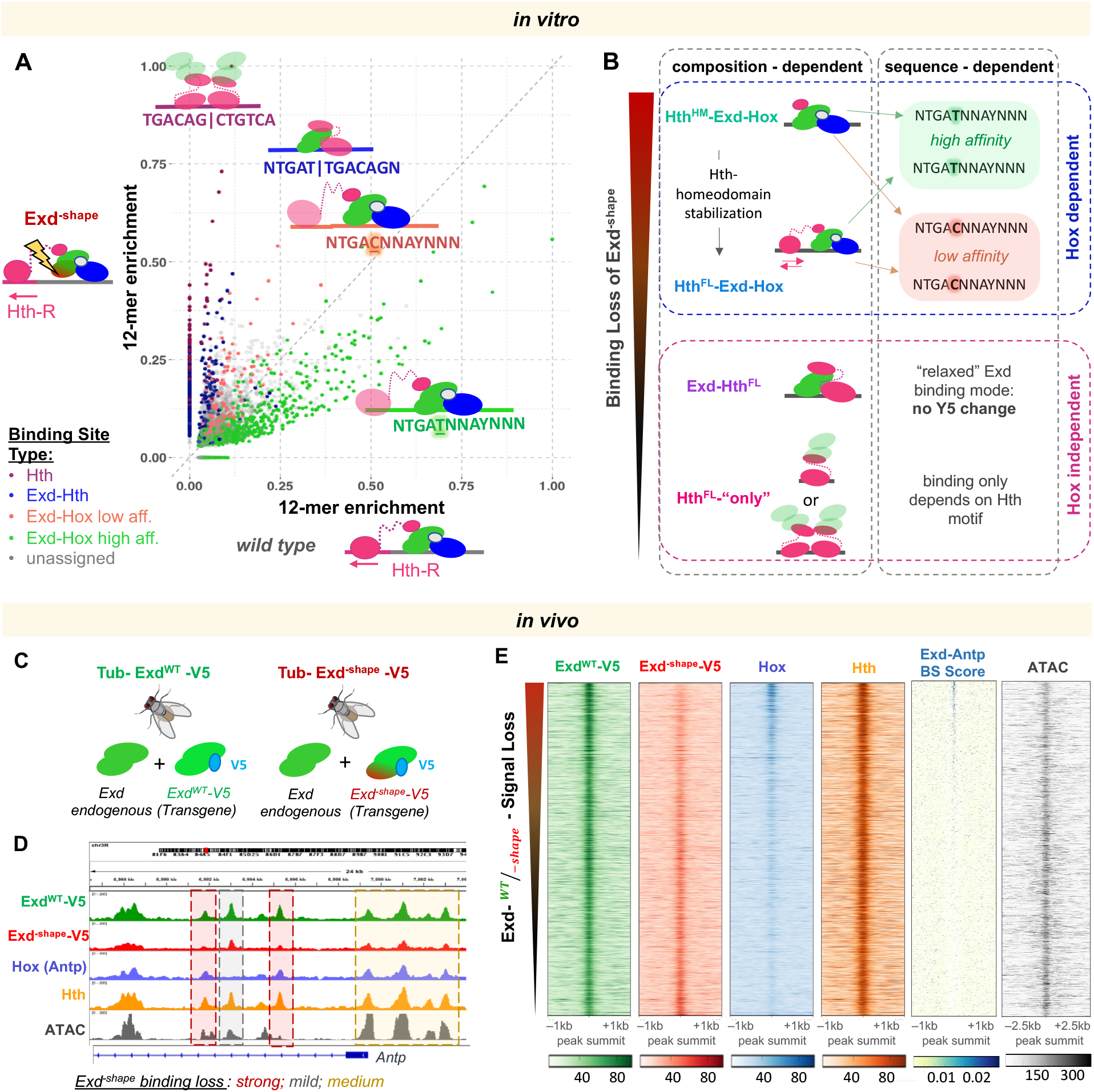
A shape readout mutant distinguishes between TF complexes *in vitro* and *in vivo*. **(A)** Classification of 12-mer DNA sequences in terms of their observed *in vitro* relative enrichment (Lib-Hth-R) in the presence of Hth^FL^-Exd-Hox and Hth^FL^-Exd^−shape^-Hox. Points/sequences are colored according to which particular HD complex best explains their enrichment: Hth dimers (purple), Exd-Hth^FL^ (dark blue), or Hth^FL^-Exd-Hox (low-affinity: Y5=C, NTGA**<u>C</u>**NNAYNNN, coral red; or high-affinity: Y5=T, NTGA**<u>T</u>**NNAYNNN; green). **(B)** Schematic illustrating the context dependence of binding loss due to the Exd^−shape^ mutation. **(C)** To perform *in vivo* validation, transgenes carrying either Exd^WT^ or Exd^−shape^ tagged with V5 were inserted into the attp40 landing site on chromosome II in the background of endogenous Exd. **(D)** Tracks showing, at the *Antp* locus, the result of anti-V5 ChIP-seq experiments performed on third instar larval wing discs of flies homozygous for *tub>exd^WT^-V5* (green) or *tub>exd*^−shape^-*V5* (red) transgenes and endogenous Exd. Hth (orange) and Hox (Antp; blue) ChIP-seq, as well as ATAC-seq (gray) tracks are also shown for reference. Background shading indicates peaks that are strongly lost (red), mostly unaffected (gray), or partially lost (yellow) by the Exd^−shape^ mutation. **(E)** Raw coverage tracks around the Exd^WT^-V5 ChIP-seq peak summit for IP signals from Exd^WT^-V5, Exd^−shape^-V5, Hox, and Hth, along with binding site (BS) affinity scores for Exd-Antp, and ATAC-seq signal. Peaks are ordered by the Exd^WT^-V5 over Exd^−shape^-V5 IP-signal ratio (“Exd^−shape^ binding loss”).

Unexpectedly, this analysis also revealed a change in sequence selectivity for the trimeric HD complex in which Exd^−shape^ participates. Two distinct classes can be distinguished: binding was less affected when the Y5 base-pair in the Exd-Hox heterodimer site (NTGA**<u>Y_5_</u>**NNAYNNN) was C-G instead of T-A (**Fig. 3A** and **Fig. S4D**). Interestingly, the base identity at Y_5_ only impacted the ternary Exd-Hox complex, as no such difference was observed when SELEX-seq was performed with Exd^−shape^ and Hth alone (**Fig. S4D**). Notably, the T5-to-C_5_ transition is predicted to widen the minor groove at the position where the Hox spacer interacts with the DNA (**Fig. S4J**). A smaller differential effect was observed at the N_1_ position (<u>**N**</u>_1_TGAYNNAYNNN) (**Fig. S4B,C,E,F,H**), which can also be explained by a change in intrinsic MGW (**Fig. S4I**), yet is not specific to the Exd-Hox subcomplex.

Taken together, these observations suggest that in parallel to optimized hydrogen-bonds with the *α*3 recognition helices, high-affinity binding sites for multi-protein TF complexes have an optimized DNA shape characterized by a set of MGW minima at specific positions. Losing the ability to interact with individual MGW minima affects the binding of some complexes more than others (**Fig. 3B**). Thus, TFs utilize distinct DNA recognition modes depending on their binding partner (**Fig. S3B**).

### Combinatorial in vitro TF-complex binding behavior is recapitulated in vivo

If *in vivo* occupancy is governed by the same binding rules and composition-dependent sequence preferences as *in vitro*, we might be able to explain more of the observed *in vivo* binding patterns by using mutant TFs tailored to lose binding free energy contributions from a specific minor groove interaction. To test this idea, we generated transgenic fly lines that ubiquitously express a V5-tagged version of Exd^WT^ or Exd^−shape^ (**Fig. 3C**; see **Methods**). Ubiquitous expression of Exd^WT^-V5, but not Exd^−shape^-V5, fully rescues an *exd* null mutant, demonstrating that the two N-terminal-arm arginines are critical for viability (**Fig. S5A**). Because the nuclear localization and therefore the activity of Exd depends on its interaction with Hth, we confirmed that nuclear import of Exd^−shape^-V5 was not compromised (**Fig. S5B**). To investigate whether lethality in Exd^−shape^-V5 is linked to a selective loss of distinct Exd-containing complexes, we carried out whole-genome ChIP-seq assays against the V5 tag of either Exd^WT^-V5 or Exd^−shape^-V5 (in the presence of endogenous Exd) in wing imaginal discs (**Fig. 3D**). We also used ChIP-seq to characterize the genome-wide binding patterns of Hth and the Hox protein Antp, which is strongly expressed in wing discs. Visual inspection of the raw IP coverage tracks for Exd^WT^-V5, Exd^−shape^-V5, Hth, and Antp at the Antp gene locus revealed that some Exd peaks are more sensitive to the N-terminal arm shape-readout mutations than others (**Fig. 3D**). Strikingly, binding signal loss (defined as the Exd^WT^-V5 over Exd^−shape^-V5 coverage ratio at each Exd^WT^ peak summit) showed a strong correlation (*r* = 0.37, *p* < 2.2×10^−16^) with predicted relative affinity for Hth^HM^-Exd-Antp (*45*), confirming that this aspect of *in vitro* binding is recapitulated *in vivo*; by contrast, ATAC-seq data from wing discs did not show an obvious correlation (**Fig. 3E**).

We next focused on the subset of Exd^WT^-V5 peaks that contain a match to the TGAYNNAY Exd-Hox consensus site (~20% or 752 peaks total). Recapitulating our *in vitro* findings, the 30% of these where occupancy is reduced the most in Exd^−shape^ are significantly more likely to have a high predicted affinity for Exd-Hox compared to the remaining 70% (*p* = 2.0×10^−9^; T-test; **Fig. S5C**). At the same time, these peaks are significantly less likely to contain strong Hth-monomer sites (*p* = 3.6×10^−3^; T-test; **Fig. S5C**). Even more strikingly, when comparing between low-affinity Y_5_=C and high-affinity Y_5_=T for sites of type NTGAY_5_NNAY, the altered sequence selectivity for Exd^−shape^ identified *in vitro* was recapitulated *in vivo*, with Antp and Exd^WT^ preferring Y_5_=T over Y_5_=C sites, with the opposite binding preference observed for Exd^−shape^ (*p* < 2.2×10^−16^ (Antp); *p* = 0.01 (Exd^−WT^); *p* = 8.2×10^−04^ (Exd^−shape^); T-test; **Fig. S5D**). That the difference between these two classes is more pronounced for the Antp profile than for the Exd^WT^ profile suggests that while Exd-Hox binding is the dominant mode, Exd-Hth complexes might compete for these same sites *in vivo* (cf. Fig. S1A), therefore contributing to the overall Exd^WT^ IP signal and reducing the effect size.

### Identification of complex composition in vivo on a genome-wide scale

Given that the stability of each type of HD complex is impacted to a different degree by the Exd^−shape^ mutation (cf. Fig. 3A,B), we reasoned that using the mutant binding loss as a diagnostic feature, along with relative affinities predicted from *in vitro* SELEX data, might allow us to classify all Exd^WT^ peaks (~3,700) in terms of a particular homeodomain complex (see **Methods**).

Using a combination of three ChIP enrichment values and three predicted binding affinity scores, each Exd peak was assigned to one of eight clusters (**Fig. 4A**). Interpretable and distinct clusters were only obtained when ChIP coverage for both Exd^−shape^-V5 and Hox was included among the features (**Fig. S5E**). Based on these data, we assigned each cluster to a particular type of complex, and whether the binding was low or high affinity (**Fig. 4A**). The resulting classification closely recapitulates that based on *in vitro* data in Fig. 3B. One cluster, comprising only 129 peaks, showed high mean values for all features, indicative of the ternary Hth^FL^-Exd-Hox complex (**Fig. 4A**).

**Fig. 4:**
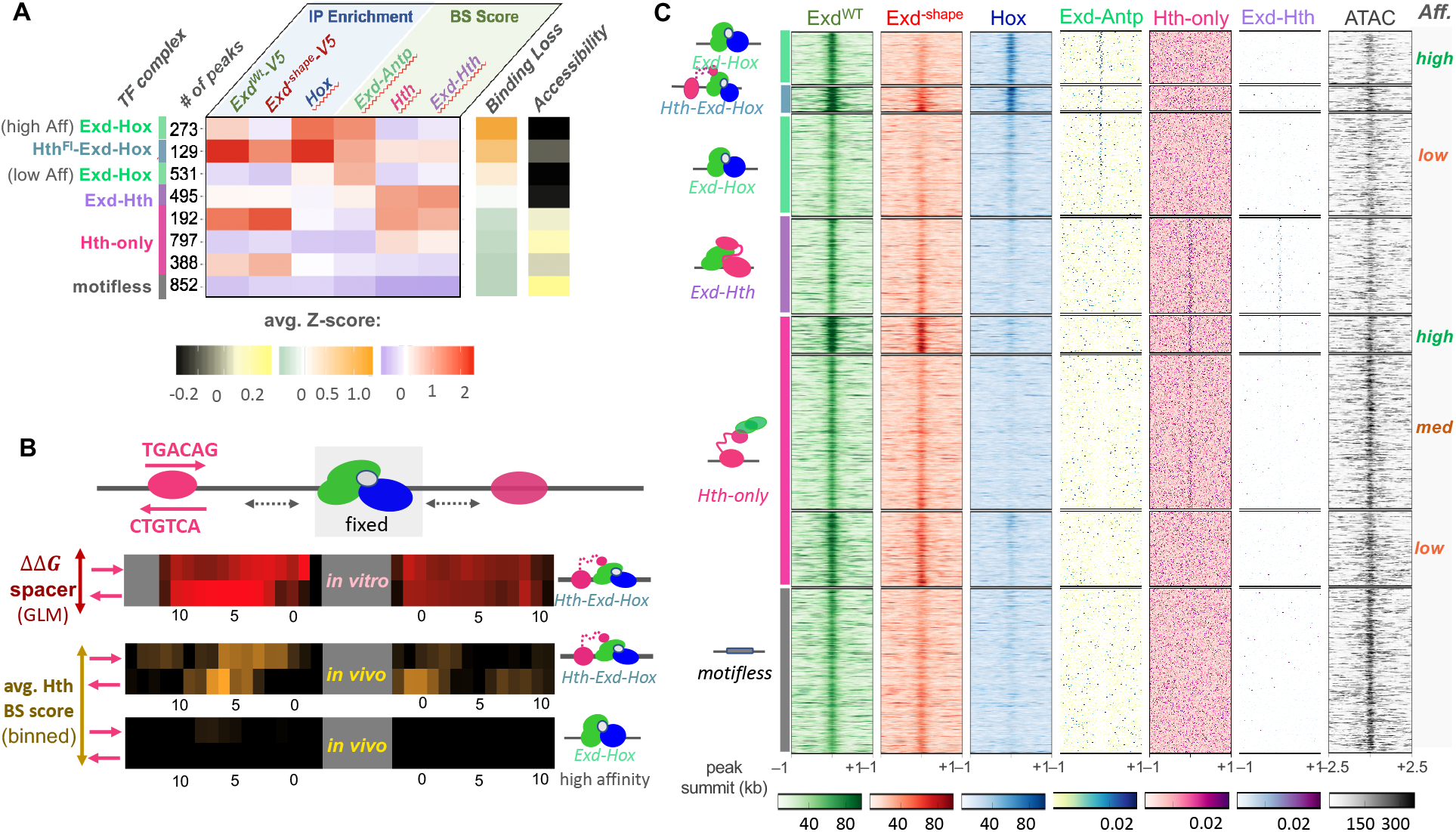
Attribution of binding complex composition *in vivo* using the Exd^−shape^ mutant. **(A)** Classification of all Exd^WT^-V5 peaks based on ChIP-seq enrichment for Exd^WT^-V5, GFP-Antp, and Exd^−shape^-V5 and predicted binding site affinity for Exd-Antp, Hth-monomer, and Exd-Hth. The heatmap shows average Z-scores across all six input features for each cluster. Average Z-scores for Exd^−shape^ binding loss and ATAC-seq signal (the latter not used for the clustering) are shown using orange-green and yellow-black color scales, respectively. The number of peaks per cluster and the assigned complex are indicated on the left. **(B)** Comparison of length preferences for the spacer between the Exd-Hox and Hth binding sites between *in vitro* and *in vivo* context. Estimated binding free energy for all four possible Hth configurations centered around the [Exd-Hox]f site derived from SELEX data is shown in the top panel (red-black color scheme). The middle and bottom panel indicate the 4-bp moving average binding site score for Hth centered around the highest-scoring Exd-Hox site for either the 129 trimer (middle) or 273 high-affinity Exd-Hox cases (bottom). **(C)** Raw tracks for IP coverage and binding affinity centered around each peak summit for all six input features, along with the ATAC-seq signal. The deduced identity of the bound complex for each cluster is indicated on the left and affinities are shown on the right.

Having identified potential trimer sites *in vivo*, we tested whether the differences in spacer preference we observed *in vitro* (cf. Fig. 2A) could also be seen *in vivo*. To this end, we aligned all 129 trimer peaks by their highest-affinity Exd-Hox site, scored Hth^FL^ binding affinity in either orientation up- and downstream of that site, and averaged over a 4-bp moving window. Indeed, distinct spatial preferences were observed, which paralleled the *in vitro* trends (**Fig. 4B**). As expected, an enhanced Hth-monomer binding affinity score was not observed for the 273 high-affinity Exd-Hox peaks (**Fig. 4B**).

Interestingly, even though ATAC-seq signal intensity was not included as a feature in the clustering, it correlated with complex composition and configuration: Sites where Exd directly contributes to DNA binding by its canonical head-to-tail Exd-HD orientation (i.e. Hth^HM^-Exd-Hox and Exd-Hth) were less accessible than sites that contain a Hth binding site bound independently of Exd (i.e. Hth-only and Hth^FL^-Exd-Hox; **Fig. 4A**). This observation suggests that different TF-complexes or configurations might have opposing effects on DNA accessibility and gene expression.

### Exd binding sites interact in 3D

To determine how many Exd peaks within each cluster of Fig. 4A can be explained by one of the three distinct binding affinity models, (Exd-Antp, Exd-Hth, or Hth-monomer), we visualized the raw input features used in our unbiased clustering (**Fig. 4C**). Strikingly, nearly 80% of all peaks showed enrichment for a complex- and cluster-specific sequence motif around the peak summit and only about 20% of the peaks remained unclassified. To investigate the molecular mechanism by which these motifless Exd peaks are established, we performed *in situ* chromatin capture (Hi-C) (*46*, *47*) on third instar larval wing discs (**Fig. 5A**). First, we asked whether Exd binding sites tend to associate in 3D space. To test this, we generated Hi-C maps at 5-kbp resolution (*48*) (see **Methods**) and extracted chromatin interaction frequencies between all pairs of Exd peaks. We found a surprising level of structure in these Exd-centric interaction maps (**Fig. 5A**). To rule out the possibility that a random sample of accessible genomic loci might produce a similar pattern, we generated a reference distribution by randomly selecting size-matched sets from all ATAC-seq peaks. Indeed, Exd peaks were about twice as likely to show significant contacts as randomly chosen sites (p-value = 4.9×10^−54^), even when only considering non-duplicated bins (p-value = 2.6×10^−32^, **Fig. S6A**). This suggests that Exd-containing chromatin-bound complexes co-localize within the nucleus. A prominent example of this is found on chromosome 2L, where many Exd peaks cluster within a region of about 200 kbp (**Fig. 5A**), containing genes such as *no ocelli* and *elbow B*, which are involved in eye-antennae development (*49*), and genes related to neuronal function such as *pickpocket* (an ion channel) and *Partner of Bursicon*, part of the Bursicon neurohormone dimer (*50*).

**Fig. 5:**
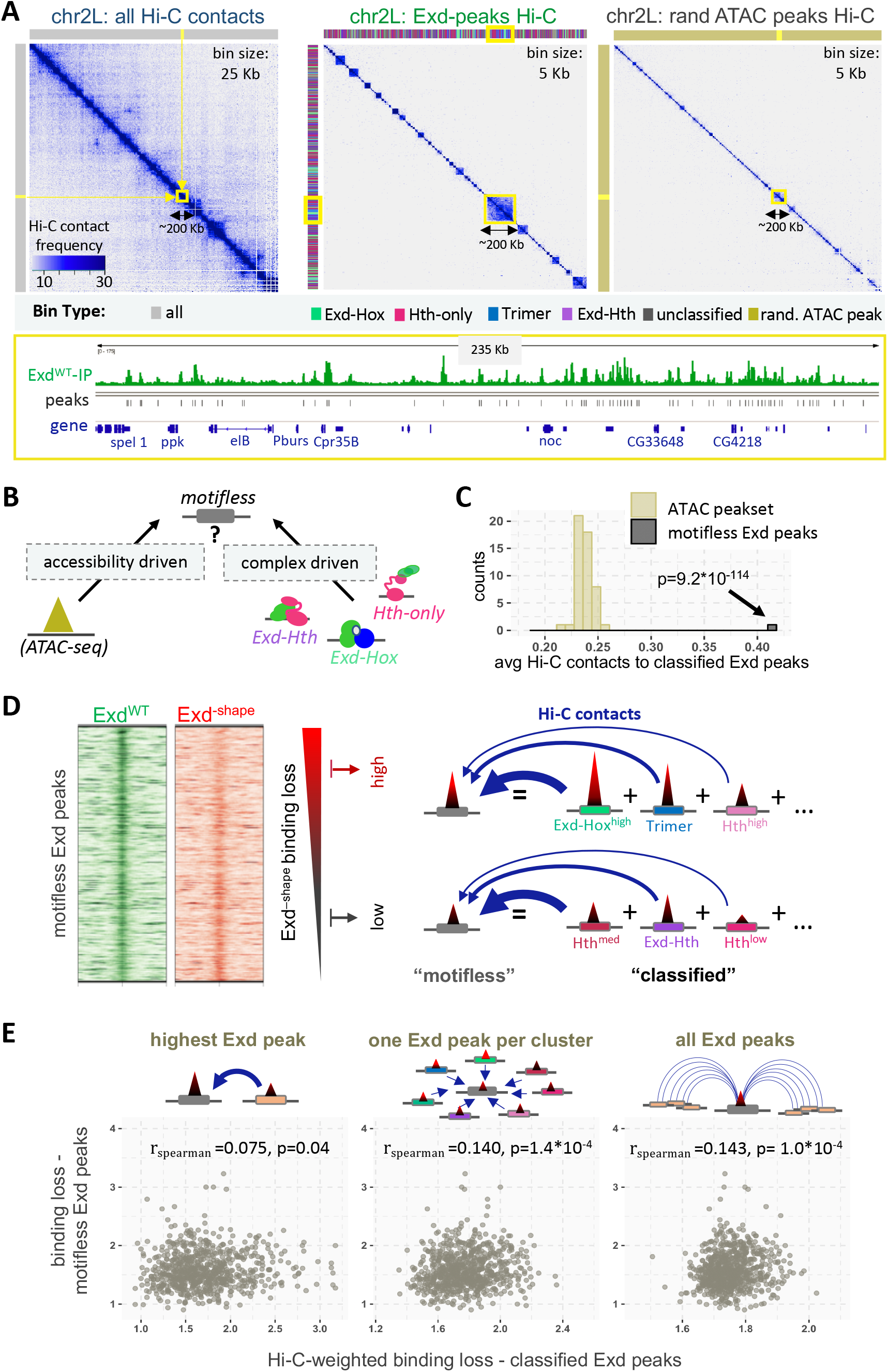
Exd binding sites form networks in 3D space. **(A)** Hi-C contact maps of wild-type (including tub>exd^WT^-V5 transgene) third instar larval wing discs for chromosome 2L showing either all chromatin contacts (left; binned at 25 Kb resolution), a selection based on the set of all Exd peaks (middle; binned at 5Kb resolution), or one based on a size-matched random sample of ATAC-seq peaks (right; binned at 5Kb resolution). Color bars above and next to each plot show the type of chromatin bin. The gene structure and raw Exd^WT^ IP signal of the highlighted area on the Hi-C maps (yellow box) is shown below. **(B)** Schematic representation of two distinct mechanistic hypotheses regarding the Exd IP signal at “motifless” genomic sites: These sites either emerge as a consequence of high local chromatin accessibility, or they occur due to chromatin interactions with motif-driven Exd peaks. **(C)** Motifless Exd peaks show high average Hi-C contact frequency with classified Exd peaks compared to a size- and accessibility-matched sample distribution of ATAC-seq peaks. **(D)** The degree of IP signal loss at motifless binding sites in response to Exd^−shape^ mutation is variable, which might be caused by 3D contacts with classified Exd binding sites. **(E)** Exd^−shape^ binding loss at motifless Exd peaks may be explainable in terms of binding loss at classified Exd binding sites it interacts with in 3D (optionally weighted by Hi-C contact frequency). Three different models are compared: (i) one that uses the unweighted Exd^−shape^ binding loss at the classified site with highest Hi-C contact frequency as a predictor; (ii) ones that takes the weighted Exd^−shape^ binding loss for one classified Exd binding site per cluster into account (cf. Figure 5; middle); or (iii) one that uses the weighted Exd^−shape^ binding loss across all classified Exd peaks located on the same chromosome.

If the 20% of Exd peaks that are not explained by a motif are merely the result of high accessibility, they should exhibit a similar degree of physical interaction with classified Exd peaks as highly accessible ATAC-seq peaks (**Fig. 5B**). Interestingly, we find the opposite to be true: Motifless Exd peaks are much more likely to be in physical proximity to sequence-specific Exd peaks than randomly sampled, highly accessible sites (p-value = 9.2×10^−114^; **Fig. 5C**). This indicates that their presence is a consequence of frequent colocalization with sequence-specific Exd binding sites, which in turn suggests that motifless Exd sites inherit their sensitivity to the Exd^−shape^ mutant from the Exd sites that they interact with (**Fig. 5D**). Supporting this idea, binding loss in the Exd^−shape^ mutant at the classifiable Exd sites, combined using the Hi-C contact frequency as a weight (see **Methods**), showed a statistically significant correlation with the binding loss at unclassified Exd sites (**Fig. 5E**). Importantly, this correlation improved when more than one contact with classified sites was considered, arguing that the IP signal at motifless sites is accrued through multiple interactions, possibly occurring within a transcriptional hub.

### The Exd^−shape^ mutant reveals distinct biological functions for different complexes

Since the Exd^−shape^ mutant predominantly impacts the binding of Exd-Hox complexes, we should in principle be able to identify the gene network directly controlled by Exd-Hox. To circumvent the lethality caused by the Exd^−shape^ mutation, we tagged the endogenous Exd C-terminally with Green Fluorescent Protein (Exd-GFP) and used the deGradFP method to deplete endogenous Exd^GFP^ protein (*51*) (**Fig. 6A and Fig. S7A**). After expressing the deGradFP system for 24 hr, we performed RNA-seq on third-instar imaginal wing discs of male flies that carried either a copy of Exd^WT^ or of Exd^−shape^ (**Fig. 6A**). At a false discovery rate of 5% we detected 392 genes upregulated in Exd^−shape^ relative to Exd^WT^, and 322 downregulated genes (**Fig. 6B**). Among the former were *exd* and *hth*, which showed mild upregulation suggestive of an autoregulatory feedback loop for Exd-containing complexes.

**Fig. 6:**
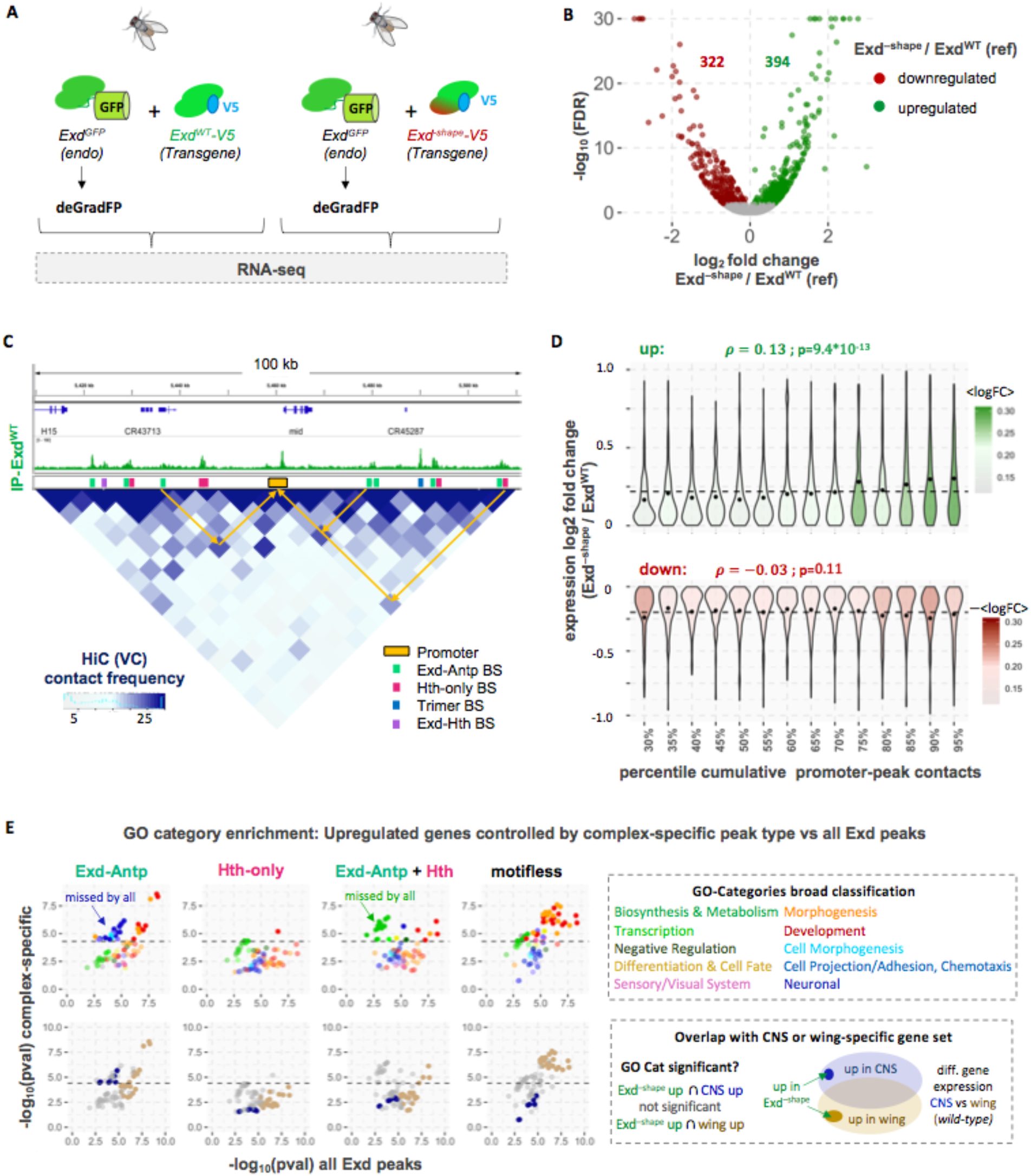
Harnessing the Exd^−shape^ mutant to decipher complex-specific biological functions. **(A)** Using the Exd^−shape^ mutation as a genetic tool to dissect the gene expression response of Exd-Hox binding loss *in vivo*. CRISPR-Cas9 based tagging of the endogenous Exd locus with GFP allows time-controlled removal of endogenous Exd protein using the deGradFP system in the background of either tub>exd^WT^-V5 or tub>exd^−shape^-V5 transgenes. **(B)** Volcano plot of the false discovery rate (FDR) versus the log_2_-expression-fold change in Exd^−shape^ compared to Exd^WT^ (reference) is shown. Genes upregulated in the Exd^−shape^ line are shown in green; downregulated genes in red. **(C)** Using Hi-C data to assign peaks to the promoter they contact the most. Shown is a region on chromosome 2L encompassing the *mid* gene locus. Exd^WT^-V5 IP coverage track is shown above the Hi-C map at 5-kbp resolution. Promoter regions (orange) and different HD complex types are shown as colored boxes. Arrows indicate examples of contacts in 3d space between enhancers (peaks) and promoters. **(D)** Cumulative promoter to Exd-peak contact frequency is significantly correlated with expression log_2_-fold-change for upregulated (green), but not downregulated genes (red). **(E)** Contact-based promoter-peak type assignment reveals distinct functions for Exd-Antp target genes. Enrichment of specific Gene Ontology (GO) categories was analyzed either based on all upregulated genes associated with any Exd peak (X-axis) or only those genes associated with a specific Exd complex in our previous analysis. Dotted lines indicate the p-value threshold at which significance is met after accounting for multiple hypothesis testing. Bottom panel shows whether the same GO categories are significantly enriched among genes both upregulated in the Exd^−shape^ mutant and specifically expressed in the central nervous system (CNS, dark blue) or wing disc (khaki) based on a transcriptome analysis comparing wild-type larval CNS and wing. “missed by all” highlights GO categories that were not identified when the entire set of Exd peaks was analyzed.

We next asked to what extent our Hi-C data, which allow us to assign Exd peaks to gene promoters, can be used to predict differential gene expression (**Fig. 6C**). We argued that the cumulative contact frequency of all Exd peaks a gene promoter interacts with within 50 kbp might be a reasonable predictor for how the expression of a given gene responds to the Exd^−shape^ mutation. Indeed, we observed a significant positive correlation between cumulative peak-promoter contact frequency and expression log-fold-change for all upregulated genes (Pearson correlation *r* = 0.13; p = 9.4*10^−13^) (**Fig. 6D**). Among downregulated genes, the same correlation was not significant (*r* = 0.03; p = 0.11). Upregulated but not downregulated genes also have significantly more contacts with all classifiable Exd peaks split by complex (**Fig. S7B**). The lack of differentiation between complexes likely results from the high degree of physical interaction among Exd peaks (cf. Fig. 5A). However, because Exd-Hox binding is most strongly affected by the Exd^−shape^ mutation, the observed changes in gene expression are likely to be driven by the loss of this particular complex. To test this idea, we again used our Hi-C data to identify the most frequently contacted promoter for each peak. Analyzing Gene Ontology (GO) associations showed that the genes directly controlled by Exd-Hox are enriched for several distinct functions that are missed when no discrimination is made among the various Exd-containing complexes (**Fig. 6E**). Among those functions were several neuronal categories, such as axon guidance, chemotaxis, and cell projection/cell morphogenesis related ones. Accordingly, the same GO categories scored significantly when taking the overlap between upregulated genes in Exd^−shape^ and genes more highly expressed in the wild type central nervous system (CNS) compared to wild type wing discs (**Fig. 6E**). As expected, no enrichment for particular GO categories was observed for the Hth-only class of Exd peaks, suggesting that they predominantly occur downstream of Exd-Hox. Only when the subset of genes controlled by both an Exd peak classified as Hth-only and one classified as Exd-Hox was considered, categories related to biosynthesis and metabolism emerged as enriched (**Fig. 6E**). Surprisingly, when analyzing the gene set associated with motifless Exd peaks we found the same, very broad functions enriched as when analyzing the gene set associated with any class of Exd peaks (**Fig. 6E**), providing further evidence that their occurrence is not purely driven by genomic accessibility.

## DISCUSSION

Accurate prediction of which DNA sequences a given TF or TF complex will bind *in vivo* is a hard and still unsolved problem, despite the availability of many complementary high-throughput datasets. There are two major reasons why we fall short of this goal: First, any one TF can bind DNA sequences with a variety of partners, such that the simplifying assumption that a single binding mode captures the full range of binding behaviors is unlikely to hold true. Second, we currently lack methods that reliably incorporate information on chromatin conformation (*52*) and therefore fail to predict sites that lack a distinct sequence signature, yet physically interact with functionally bound sites.

In this study we showcase how we can infer structural features of multi-TF complexes from high-throughput *in vitro* binding data. Importantly, the mechanistic insights we obtain challenge several currently held views on the nature of TF binding, including the subordinate role that structurally ill-defined protein regions and DNA sequences lacking readily defined motifs play in TF-target recognition and complex stability. Our biophysically motivated but otherwise unbiased analysis of SELEX-seq data allowed us to specifically design TFs that can be deployed as tools to interrogate complex TF binding behavior and function, both *in vitro* and *in vivo*. Specifically, our thorough investigation of HD binding mechanisms revealed that Exd relies on shape-readout in a complex-specific manner. In general terms, a single TF can utilize its ability to read a narrow minor groove to different degrees, depending on both complex composition and DNA sequence identity. Because quantitative information about any readout mechanism that contributes to DNA sequence specificity is implicitly captured in high-throughput binding assays, it is likely that our approach can be used to infer structural mechanisms and molecular configurations for other TF systems as well. Instead of directly generating libraries of mutant TFs and assaying their binding profile, naturally occurring protein sequence variation among TF paralogs or binding partners can be leveraged.

Our realization that a part of a TF that samples many configurations can differentially affect the assembly of distinct TF complexes *in vitro* allowed us to infer the bound complex for ~80% of all Exd peaks, a vast improvement over what can be achieved using just a single static motif. In the current literature, the remaining 20% of sites, which lack a binding motif but display high accessibility, would typically be considered a byproduct of ChIP-seq experiments (*17*, *18*). However, the key observation that the loss of ChIP-seq signal in the Exd^−shape^ mutant at motifless sites correlates with that at motif-dependent sites suggest that, at least for Exd, motifless sites inherit their binding loss from 3D interactions with sites bound by specific Exd-containing complexes. Thus, we propose to refer to motifless yet TF-bound complexes as “shadow peaks”. Notably, genes whose promoters are contacted by the subset of Exd shadow peaks show statistically significant enrichment for the same Gene Ontology categories as the entire set of Exd peaks. This suggests that shadow peaks do not reflect non-functional binding, but may mark, or even play an active role in, the formation of functionally relevant chromatin hubs.

We also made two key observations regarding combinatorial TF gene control. First, we showed that complex-specific function might be missed when analyzing all TF peaks together (cf. Fig. 6E). For Exd-Hox, we found a repressive role that limits the expression of genes biased for nervous system expression. This finding is consistent with previous observations showing that although the Hox gene *Antp* is dispensable for wing formation, removing its activity often results in morphological abnormalities (*53*). We also identified a common gene set controlled by Exd-Antp and Hth-only complexes – genes associated with metabolic function and biosynthesis. This finding may be relevant to the further investigation of the seemingly contradicting roles that Hox proteins and their cofactors play in the onset of cancer (*54*). Second, we found that the interaction frequency between Exd-bound regulatory elements and their target promoters is a quantitative predictor of differential gene expression. The correlation is significant despite downstream effects, potential redundancies among TFs, and the use of whole-tissue data rather than that of isolated cell populations, and is consistent with a recent study showing that interactions between promoters and their known enhancers were correlated with nascent RNA levels using high resolution imaging (*55*).

Lastly, we might ask to what extent these mechanisms apply to mammals: The human genome encodes four highly conserved orthologs of Exd, namely Pbx1-4 (*23*). In the mouse, where knockouts have been studied, all four *pbx* genes are essential for viability (*27*, *56*–*58*). Consequently, a complete loss-of-function (null) allele of *pbx* would be unlikely to contribute to human disease unless the gene was haplo-insufficient for a specific function. In contrast, a subtler perturbation of Pbx activity, analogous to the shape-defective mutation of Exd described here, could in principle contribute to human disease by interfering with the binding of specific Pbx-containing TF complexes. With this in mind, we examined several human genetics databases. Notably, missense mutations in Pbx1-3 homeodomains are underrepresented in healthy populations [http://gnomad.broadinstitute.org; (*59*)], consistent with the essential function of these DBDs. An interesting exception are *de novo* mutations of N-terminal arm arginines of Pbx1 that are present in several patients diagnosed with congenital anomalies of the kidney and urinary tract syndrome (CAKUTHED; (https://www.omim.org/) (*60*, *61*): Three patients had a mutation in either the R2 (1x) or R3 (2x) arginines of Pbx1, equivalent to the ones mutated here in Exd. These Pbx1 mutants were defective in their ability to activate a reporter gene harboring a perfect Exd-Hth binding site (*61*). We speculate that these human *pbx1* alleles are essentially DNA shape readout defective mutants of Pbx1 and, as a result, are compromised in the binding of a particular subset of Pbx1-containing TF complexes to their respective binding sites, resulting in the highly specific CAKUTHED syndrome.

## Acknowledgments

We thank the members of the Lomvardas lab for sharing their experience in performing Hi-C experiments, and Remo Rohs, as well as the members of the Bussemaker, Mann, and Rohs labs, for valuable discussions.

## Author contributions

This J.F.K., R.S.M., and H.J.B. designed the study and conceived of the experiments; J.F.K. carried out the experiments with help from R.E.L., S.F., and N.A.; J.F.K. and H.J.B. constructed the feature-based models with input from H.T.R.; J.F.K. performed the data analysis; C.R. generated the NRLB models; J.F.K., R.S.M., and H.J.B. interpreted the data and wrote the paper. All authors discussed the findings and contributed to the manuscript.

## Data and materials availability

The raw sequencing data for both *in vitro* (SELEX-seq) and *in vivo* (ChIP-seq, ATAC-seq, RNA-seq and Hi-C) experiments as well as the Exd peak set and coverage tracks used for downstream analyses will be made available through the NCBI Gene Expression Omnibus.

## Materials and Methods

### Protein purification and mutagenesis

Fly proteins were obtained and purified as described in (Slattery, 2011). Briefly, PET-expression vectors containing coding regions for full-length hth (Uniprot-ID: O46339), exd (Uniprot-ID: P40427), dfd (Uniprot-ID: P07548) and Hth HM-domain (amino acids 1-242; (Uniprot-ID: O46339) with hexa-histidine tags (except for Exd, which was copurified with full-length Hth or HM-domain-only Hth) were transformed into Bl21 cells. Cells were grown for 5-7 hours, lysed, and proteins extracted with affinity purification using Cobalt-Talon beads (Clontech). Site-directed mutagenesis for Exd and Hth was performed via amplification of the original plasmid with primers harboring single amino acid replacements (arginine to alanine) using Taq-polymerase (NEB). Double and triple mutations were generated consecutively. **Supplemental Table S1** contains a summary of the mutations made.

### Binding and competition assays

Electromobility shift assays (EMSAs) were performed using 2 nM radiolabeled DNA and protein concentration between 75-900 nM: Dfd was kept constant at 150 nM; wild-type Hth^FL^-Exd and Hth^HM^-Exd was used at 100 nM; mutant proteins were increased from 75 nM – 300 nM (two lanes) or up to 900 nM (three lanes). Proteins were incubated for at least 30 min prior to loading in binding buffer (final concentration: 2% Glycerol, 30 *μg/μl* polydIdC, 40 mM NaCl, 40 mM Tris pH=8.0, 0.4 mM MgCl2, 1mM DTT, 0.5 mM EDTA). After loading onto a 5% TBE gel, gels were run at 4°C for 2h in 0.5x Tris-running buffer. For competition assays, a radio-labeled probe was competed out with increasing concentrations of unlabeled competitor DNA while keeping protein concentrations constant (100 nM). Dose-response curves and IC50 values were obtained using the R package drc. Spacers with zero, three and seven bases between the Hth and Exd-Dfd sites were tested.

### Library design

The Lib-16 library contained a 16-mer random flank without fixed binding sites and data for Hth^HM^-Exd-Dfd were taken from Slattery et al. (*1*). The data for the Hth^FL^-Exd SELEX-experiment was generated using a Lib-16 library as well, but following the design described in (*2*). The Lib-Hth-F and Lib-Hth-R libraries contained a fixed Hth site – TGACAG in forward (F) and CTGTCA in reverse (R) orientation – immediately followed by a 21 bp random region. Library Lib-30 had a 30-bp random region and no fixed Hth binding site. Full library sequences are listed in Supplemental Table ST2.

### SELEX experiments

For Lib-Hth-F, Lib-Hth-R, and Lib-30, SELEX experiments were carried out using wild-type or mutant homeodomain proteins following the experimental procedures described in (*1*, *3*). Two rounds of enrichment were performed for each set of experiments. For the Hth^FL^-Exd and Hth^Fl^-Exd^−shape^ SELEX-experiments using Lib-16, a single round of selection was performed using the methodology and library design described in (*2*). Data for Hth^HM^-Exd-Hox was obtained from a previous study (Slattery, 2011). For each experiment, proteins of a final concentration of ~50 nM were assembled and incubated with excess DNA (10-20 fold) for 30 minutes. After each round of selection, the DNA was extracted from the gel amplified by either using Ilumina’s small RNA primer sets or the set of primers described in (*2*). Sequencing barcodes were added in a five cycle PCR step and the final library was gel-purified using a native TBE-gel before sequencing.

### Sequencing and data processing

Libraries for Hth^FL^-Exd and Hth^FL^-Exd^R2A,R5A^ (Lib-16) were sequenced using a v2 75-cycle high-output kit on an Illumina NEXTSeq Series desktop sequencer at the Genome Center at Columbia University. Libraries Lib-Hth-F and Lib-Hth-R with either Hth or Exd shape-readout mutant in complex with the respective other wild-type protein and Dfd, as well as the Lib-30 Hth^FL^-Exd-Dfd experiment were all sequenced at the New York Genome Center using separate lanes on an Illumina HiSeq 2000 sequencing machine. Libraries Lib-Hth-F and Lib-Hth-R with wild-type proteins were also sequenced on a HiSEQ instrument at a different facility. Libraries were trimmed to remove Illumina- and library-internal adapter sequences using custom shell scripts and the FASTX toolkit (Hanon lab) and loaded into the R environment using the R package named SELEX (http://bioconductor.org/packages/SELEX) (Riley, 2014).

### Computational analysis of complex composition and orientation

Relative enrichment tables for all libraries were generated using the SELEX package. To color the individual oligomers based on the complex composition most likely explaining their enrichment, position-specific-affinity matrices were generated for Hth^HM^-Exd-Dfd using a 12-mer seed sequence from Lib-16 and Lib-Hth-(F/R) (12-mer with highest enrichment), for Hth^FL^-Exd using a 10-mer or 12-mer seed (TGATTGACAG or TTGATTGACAGC), and for dimeric Hth^FL^ using a 12-mer seed (TGACAGCTGTCA; Lib-30). Each sequence from each respective library was then scored with the different PSAMs and complex composition assigned based on the PSAM achieving the highest score. To remove shifted binding sites that do not encompass the full TF footprint, only sequences with a relative affinity score or > 0.01 for one of the three PSAMs were retained.

To test for preferences in complex orientation with respect to the fixed Hth site in the Lib-Hth-(F/R) libraries, overall 12-mer relative enrichment tables were generated as described above and forward or reverse-complement orientation assigned by comparing the relative enrichment of each 12-mer to that of its reverse complement. Sequences with a higher score for the forward strand (as obtained from the sequencing run) were designated as [Exd-Hox]_F_ and sequences with a higher score for their reverse complement as [Exd-Hox]_R_. Average F/R ratios for Lib-16 (Hth^HM^-Exd-Dfd) and Lib-Hth(F/R) were shown as boxplots (**Fig. S1C**). To account for different offsets of the Exd-Hox complex, 12-mer enrichment tables were generated for each offset respectively (using the SELEX function selex.affinities(…, offset=x) with x=0 to 9) and F and R orientation assigned accordingly. To test for sequence preferences within the DNA spacer connecting the Hth and Exd binding sites, 16-mer enrichments of sequences right downstream of the fixed Hth site of Lib-Hth(F/R) (offset = 0) were computed and sequences isolated that matched the top Exd-Hox binding site (ATGATTAATGAC) at position 5-16. A+T content of the variable 4bp spacer sequence was computed and compared to the relative enrichment of each 16-mer (spacer) sequence.

For the comparison of k-mer based relative enrichment plots between wild-type and shape-readout-mutant SELEX libraries, each sequence was assigned an F or R orientation as described above, as well as a representative complex that best explained the sequence signature (using PSAMs, see above). In addition, Exd-Hox type sequences were split based on the Y_5_ (C or T) or the N1 (A,C,G,T) base identity within the consensus 12-mer binding sites (**<u>N</u>**TGA**<u>Y</u>**NNAYNNN; **Fig. 3A** and **Fig. S4**). For representation purposes, only sequences in F orientation and with a PSAM score greater than 0.005 were shown. Sequences that had similar scores (less than 3-fold difference) for more than one PSAM, or that did not match the respective Y5 pattern (e.g. due to a partial motif), were labeled ambiguous and colored separately (grey).

### Feature-based modeling using GLM

To model the relative orientation and offset preferences for the Exd-Hox subcomplex quantitatively in a unified model, each 21-bp probe sequence (including 2 bp of flanking sequence) was first scored on both strands with a PSAM obtained from the Hth^HM^-Exd-Dfd data set from Lib-16. Only probes where a unique binding site solely accounted for >95% (trimeric complex) or >90% (dimeric complexes) of the probe selection were retained. A similar procedure was described in (*4*). Probes with identical 12-mer Exd-Hox sequences, spacer length and strandedness were collapsed to one entry in the design matrix. The collapsed R2 counts were used as dependent variables in the generalized linear model, log-transformed respective R1 counts were used as an offset and both log-transformed Lib-16 derived relative enrichments for the Exd-Hox subcomplex and the overall configuration, as defined by the combination of spacer length and the orientation of Exd-Hox ([Exd-Hox]_F_ or [Exd-Hox]_R_) were used as predictors/features in the model. The model was fit using the R function glm(…, family=poisson) based on the following model, where S_i_ represents the sequence of the Exd-Hox 12-mer with a specific configuration:

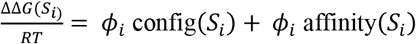

Oligomer-based models for sequence preferences within the spacer were obtained using the same modeling framework. The full set of confidence-filtered probes was first subsetted by offset (spacer length) and orientation. Choosing a specific offset *L* (e.g. spacer of length *L*=4) and Hth-[Exd-Hox]_R_ orientation, sequences identical over *L*+12 bases where first collapsed and the total R2 occurrence was used as the response variable in the model. The log-transformed Markov model predictions for the R0 initial bias of each (*L*+12)-mer was used as an offset and the spacer sequence and the relative enrichment value for each 12-mer, were used as predictors, resulting in 4^*L*^ + 1 model predictors.

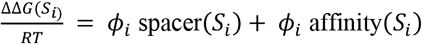

For the mononucleotide model, the oligomers were represented by 4*(*L*+12) base identity indicators, reducing the parameter space:

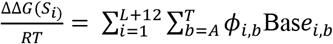

Model comparisons were done by computing the R^2^ (based on a linear model) between the spacer coefficients from the oligomer model and the sum of the base coefficient making up the respective spacer sequence in the mononucleotide model (**Fig. S1I**).

Models with fixed N_1_N_2_ base identity were obtained by further subsetting the probes, such that the Exd-Hox binding site would start with AT (see optimal 12-mer sequence). Fixing the first two positions allows isolating shape-dependent sequence selection within the spacer from effects due to readout occurring within the core Exd-Hox binding site. Mononucleotide models were fit for different spacer lengths as described above, while excluding the first two base positions within the Exd-Hox site from the feature set.

### Affinity-shape correlation

To identify whether shape might be responsible for the observed spacer selection, we first computed the theoretical model score ΔΔ*G*(spacer)/RT for each possible spacer, by summing up the respective base coefficients:

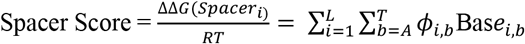

With a score for each spacer in hand, we next used the pentamer shape table (*5*) to compute the predicted minor groove width for each spacer. Since the score for each base in the pentamer table is dependent on the two bases up- and downstream, we extended the spacer 5’ with the fixed Hth binding site, present in Lib-Hth-(F/R), and 3 ‘ by the base identity of the fixed N_1_N_2_ = AT used in the model. The resulting MGW profiles for each spacer were ranked by their ΔΔ*G*(spacer)/*RT* and average MGW profiles were obtained by taking the position-wise average across sets of spacer sequences. To test for a role of MGW in selection, we first computed the average MGW profile including all spacers, setting a reference point of random or no selection. We then subsequently increased the threshold for spacers included in the analysis based on their ΔΔ*G*(spacer)/*RT* ranking and recomputed the average MGW profile on the reduced set. Sequentially removing “bad” spacers from the pool, should reveal any apparent selection for a specific MGW profile, as it mimics the underlying, biophysical selection process. Since no meaningful flank is present for the Hth^HM^-Exd-Hox and Exd-Hth^FL^ complexes, mononucleotide feature models were also obtained from the R2 or R1 counts of sequences with the core binding site extended by 4bp up- or downstream (Fig. 2D,E and Fig. S3).

### Structural interpretation

Structural representations (superimpositions) were obtained with the align function in pymol, using either the DNA (Fig.2 and Fig. S3B) as the template. Extended B-DNA with sequences accommodating the respective homeodomains and spacers were generated with the Nucleic Acid Builder webserver (*6*) (http://structure.usc.edu/make-na/server.html).

### Generation of transgenic and CRISPR-Cas9 fly lines

The full-length cDNA sequence for either the wild type or the R2A,R5A mutant Exd (obtained by PCR from the protein-expression vectors), followed 3’ (C-terminally to the protein) by the sequence coding for the small V5 peptide, was ligated into the multiple cloning site (MSC) of a vector with attB sites for *ϕ*C31-mediated integration. The vector contained a tubulin (Tub) promoter and a poly-adenylation signal surrounding the MSC. Purified vectors were sent for injection into the attp40 site on chromosome 2L, additionally marked with w+. The resulting flies were crossed with respective balancer males or females (sp/CyO; MKRS/TM2) and transgenes were tested for their ability to rescue an *exd* null allele.

For Antp ChIP-seq experiments, a GFP-tag was fused in frame into the endogenous *Antp* locus at its N-terminus (details upon request; Feng et al., in preparation), resulting in homozygous viable GFP-Antp flies.

Fly lines carrying endogenous Exd with a C-terminal Green Fluorescent Protein (GFP) tag were ordered from Rainbow, using their CRISPR-based protein tagging service. The final line harbors a GFP directly fused to the last coding amino acid of Exd, followed by an SV40-poly(A) signal and a DsRed Express cassette for easy screening. Progeny obtained from the initial red fluorescent screen were homozygous viable. Fly lines used for RNA-seq experiments were the result of a cross between i) female flies homozygous for both Exd-GFP (X chromosome) and a temperature-sensitive Tub-GAL80^ts^-UAS-deGradFP (*7*) (2^nd^ chromosome) and ii) male flies carrying either Tub-Exd^WT^-V5 or Tub-Exd^−shape^-V5 transgene on the second and an enhancer-trap into the headcase locus driving Gal4 (hdc-G4) on the third chromosome over *C(2L;3R),Tb*. Flies selected for RNA-seq were males of the following genotype: Exd-GFP/Y; Tub-Gal80^ts^-UAS-DeGrad/Tub-Exd^WT or -shape^-V5; hdc-G4/+.

### Immunohistochemistry

The following antibodies for immunohistochemistry were used: rabbit anti-Exd (*8*), mouse anti-V5 (Invitrogen, R960-25), guinea-pig anti-Hth (*9*), mouse anti-Antp (DDHB. C811), rabbit anti-GFP (Invitrogen, A-11122). Imaginal wing discs were collected from third instar larva, fixed in 4% formaldehyde for 25 minutes and stained with the antibody overnight in a 1:500 dilution. Discs were imaged at 20x magnification using confocal microscopy and processed using ImageJ software.

### ChIP-seq

The following antibodies were used in ChIP-seq experiments: mouse anti-V5 (Invitrogen, R960-25), rabbit anti-GFP (Invitrogen A-11122) for Antp-GFP, guinea-pig anti-Hth (raised against the N-terminus of Hth; GP52)(*9*). About ~ 100 third instar larval wing discs were used for each ChIP-seq sample. All buffers contained protease inhibitor (cOmplete, Roche). Inverted larvae were cross-linked at room temperature (RT) for 10 min in 10 ml 1% formaldehyde solution buffered with 50mM HEPES (pH=8.0), immediately quenched with 1 ml 2.5M Glycine and washed for 5 minutes in quench-solution (125 mM glycine, in 1X PBS and 0.01% Triton X-100). Inverted and cross-linked larvae were washed twice with Buffer A (10mM HEPES, pH=8.0; 10mM EDTA, pH=8.0, 0.5mM EGTA, pH=8.0; 0.025 % Triton-X) and twice with Buffer B (10mM HEPES, pH=8.0; 200mM NaCl, 1mM EDTA, pH=8.0; 0.5mM EGTA, 0.01 *%* Triton X-100). Wing discs were detached on ice in Buffer B and transferred into a final volume of 1 ml Buffer C (10mM HEPES, pH=8.0;1mM EDTA, pH=8.0; 0.5mM EGTA, pH=8.0). Chromatin was sheared into fragments by using a probe sonicator at 15 % amplitude (total time: 12 min with 15 seconds on and 40 second off intervals) and flash-frozen in liquid nitrogen for storage at −80°C until further processing (no more than one week). Sheared chromatin was diluted in 5X RIPA dilution buffer (1x RIPA: 140mM NaCl; 10mM HEPES, pH=8.0; 1mM EDTA, pH=8.0; 1 % Glycerol; 1% Triton X-100; 0.1% DOC) and blocked with 10*μ*g of the respective IgG-coated magnetic beads (Dynabeads, ThermoFisher) for 1h at 4°C. Beads were removed with a magnetic stand and supernatant was transferred into a new, low-binding tube. At this point, 10 % of the sample was set aside to serve as an input control. Specific antibody (10 *μ*g for mouse anti-V5, 8*μ*g for rabbit anti-GFP and 3-4 *μ*g for the Hth antibody) and 1% of Bovine Serum Albumine (BSA) was added to the remaining chromatin and incubated overnight (o/n) at 4°C. The next day, ~30 **μ*g* of IgG-coated and pre-blocked (with 1 % BSA) Dynabeads were added to each chromatin antibody solution and incubated for another 2 hours at 4°C. Antibody-bound TF-chromatin complexes were isolated by magnetic separation (5 min on a magnetic stand) and beads were washed twice with 1x RIPA, once with high salt RIPA (500mM NaCl), once with LiCl-Buffer and once with TE (10 mM Tris-Base, pH=8.0; 1mM EDTA, pH=8.0). Bead-bound chromatin and the input sample were redissolved in 0.5 ml Elution-Buffer (TE with 0.5 % Sodium Dodecyl Sulfate (SDS) and 50mM NaCl) and incubated for 30 min at 37°C with RNase, followed by 2 hours at 55°C with proteinase K (ThermoFisher). Remaining DNA-protein complexes were decrosslinked by incubating for 16 hours at 65°C. DNA was separated from the Dynabeads by magnetic separation and purified by phenol:chloroform extraction and DNA precipitation using 1x volume of isopropanol in 100 mM ammonium acetate and adding 1 *μ*l glycogen. Precipitated DNA was redissolved in 30 *μ*l TE.

### ATAC-seq

Wing imaginal discs of third instar larvae were dissected from a lab stock of *yw* genotype in Phosphate-Buffered-Saline. Discs were washed in nuclear extraction buffer (NEB, 10nM HEPES pH. 7.5, 2.5mM MgCL_2_, 10mM KCl) and placed in a 1mL dounce homogenizer (Wheaton) on ice. Discs were treated with 15 strokes of the loose pestle, followed by a 10 minute incubation on ice, then 20 strokes of the tight pestle. Nuclei were counted using a hemocytometer, and 50,000 nuclei were transferred to a fresh Eppendorf containing 1mL of NEB buffer +0.1% tween-20. Following a brief mixing the nuclei were immediately pelleted for 10 min at a speed of 1000xg. The pellet was resuspended in ATAC transposition buffer as in (*10*) and tagmentation was carried out as previously described (*10*). Amplified libraries were purified, and size-selected using double-sided ampureXP (Beckman) size selection.

### In Situ Hi-C

Wing imaginal discs of third instar larvae homozygous for both endogenous and tub>exd^WT^-V5 were dissected in PBS (with 0.5% Bovine Serum Albumin (BSA)). Discs were transferred to 1x Schneider’s Drosophila medium (Gibco) and pelleted at 300g. A single-cell suspension was generated by incubating the discs for 15 min at RT in 200 *μ*l of Schneider’s medium containing 1 *μ*g/ml of papain enzyme. The dissociation reaction was quenched by adding 800 *μ*l of Schneider’s medium with 10% Fetal Bovine Serum (FBS) and pipetting up and down at least 10 times. The cell suspension was pelleted at 600g (5 min at 4°C). Immediately after the dissociation, cells were cross-linked for 10 min (RT) in 1% methanol-free formaldehyde solution. For all subsequent steps, the protocol described in (*11*) was followed using the restriction enzyme DpnII.

### RNA-seq

Crosses to obtain flies with transgenic Exd being the dominant source of Exd were set up as described above and raised at 18°C. 24 hours before RNA isolation, larvae were shifted to 29°C. 2-4 wing discs of third instar, male, wandering, non-Tb larvae were obtained for each deGradFP RNA-seq experiment. For wild-type, larval central nervous system and wing disc RNA-seq samples, flies were raised at 25°C and 4-5 third instar wandering larvae were used per sample (3 replicates each). Discs were dissected on ice in BPS with 0.5% BSA and transferred to 350 *μ*l of RLT buffer (Qiagen) with 1% *β*-mercaptoethanol (BME). Discs were homogenized with a plastic pestle and frozen at −20°C (no more than 1 week). To each sample 100 *μ*l PBS and 250 *μ*l Ethanol was added and RNA was purified using the Qiagen RNeasy mini kit (Qiagen 74104). RNA was next treated with DNaseI (NEB) for 30 min at 37 °C, followed by another column purification using Qiagen’s RNeasy micro kit (Qiagen 74004). RNA quality was assessed with a RNA Pico Chip (Agilent) on a Bioanalyzer and only non-degraded samples were used for subsequent library generation. RNA-seq libraries were prepared using NEB’s NEBNext Ultra II Directional RNA Library Prep Kit for Illumina Sequencing (NEB EE7760S) and following the instructions for the poly(A) mRNA magnetic Isolation Module. AMPure XP beads (Beckman Coulter) were used for DNA library size-selection. DNA library quality was assessed with a High Sensitivity DNA ChIP (Agilent) on a Bioanalyzer and quantification was performed using a Qubit fluorometer. Two replicates were obtained for the Exd^−shape^ experiments and three replicates each for the CNS and wing-disc RNA-seq samples.

### ChIP-seq Library Preparation and Sequencing

ChIP-seq libraries were constructed using the NEBNext Ultra DNA Library Prep Kit for Illumina with NEBNext Mulitplex Oligos (one separate index per sample) following standard instructions. For the PCR amplification, 13-15 cycles were used depending on the amount of starting material, which was generally between 3-10 ng of precipitated DNA. For the input samples no more than 10 ng of DNA was used to match input and IP samples as closely as possible. For the final size selection, AMPure xp beads (Agencourt) were used and larger (>550bp) and smaller (<150bp) fragments were removed by a double-sided size selection with first 0.6x volume of beads to DNA and retaining the supernatant, followed by a final concentration of 0.9x beads to DNA and retaining the DNA-bound to the beads. Quality control was done by assessing the DNA size distribution with a Bioanalyzer. ChIP-seq, ATAC-seq, RNA-seq and Hi-C libraries were diluted to 2 nM, using a Qubit to verify the final concentration, pooled and sequenced with a v2 75 or a 150 cycle high-output kit using either single-end (ChIP-seq, ATAC-seq) or paired-end (RNA-seq, HiC) settings on an Illumina NEXTSeq Series desktop sequencer at Columbia University.

### ChIP-seq and ATAC-seq data processing

The four separate, raw fastq-files (from the four lanes of the sequencing run) were first collapsed into one file and subsequently aligned (bowtie2)(*12*) to the *D. melanogaster* genome version dm6 (2014, GenBank accession: GCA_000001215.4). Aligned sam files were next converted into bam files, sorted and cleared from duplicate reads using the samtools functions view, sort and rmdup (*13*–*15*). The sorted, unique bam files were indexed and converted into bigwig files using the bamCoverage function in the Deeptools suite with parameters -bs 1 -e 125 (*16*). For ChiP-seq, peaks were called using the MACS2 (*17*) function callpeak using the input samples as control files with parameters -g dm -q 0.01 or 0.05 --nomodel --extsize 125. For further downstream analysis, peak summits from the more deeply sequenced Exd^WT^-V5 ChIP replicate with a q-value threshold of 0.01 were used.

### Hi-C data processing

The four separate, raw fastq-files were first collapsed into one file. For downstream data processing the Juicer Tools Version 1.76 pipeline5 was used (*18*).The DpnII restriction site file was generated using the Drosophila genome version dm6 and the python script provided by Juicer Tools. The highest resolution to create the .hic file was set to 5 Kb. To remove multi-mappers, only reads meeting the MAPQ>30 cutoff were used. For this study only intrachromosomal contacts were considered. Contacts were dumped using the contact extraction tool Straw (*18*) with normalization method VC (vanilla coverage) at 25 Kb and 5Kb resolution. Binned Hi-C contacts were loaded into R for further data analysis.

### RNA-seq data processing and analysis

The four separate, raw fastq-files (from the four lanes of the sequencing run) were first collapsed into one file and subsequently aligned with hisat2 (*19*) to the *D. melanogaster* genome version dm6 (2014, GenBank accession: GCA_000001215.4). To obtain information on preferential promoter usage (across different isoforms), the RNA-seq data were also aligned to the most recent transcript assembly (ENSEMBL) using the program Salmon (*20*). Differential gene expression was analyzed in R using packages Rsubread (*21*) and DESeq2 (*22*). Only genes with at least 50 counts in either Exd^WT^ or Exd^−shape^ sample were used (total of ~ 8500 genes). Volcano plots were generated by using a false discovery rate (FDR) of 5% for differentially expressed genes and using the DESeq2 empirical bayes shrinkage method for fold-change estimation (*22*). For the association of contact frequency and fold change expression the same method was used.

### Coverage Plots and Downstream Peak Analysis

Heatmaps for the raw IP coverage of the four ChIP-seq samples and ATAC-seq sample (Exd^R2A,R5A^-V5, Exd^WT^-V5, Antp-GFP, Hth, ATAC-seq) were generated on the Exd peak set sorted by the Exd^WT^-V5/Exd^−shape^-V5 IP-ratio using the Deeptools functions computeMatrix and plotHeatmap (parameters: --sortRegions “no” --refPointLabel -- missingDataColor 1). Raw read coverage was extracted at the Exd peak summits (-q-value = 0.01) from the bigwig files for all ChIP samples. Further comparisons between Exd^WT^-V5 and Exd^−shape^-V5 were based on the combined coverage of both replicates. For each Exd peak, sequences surrounding the peak summit (±50bp) were extracted. Each peak sequence was then scanned with i) an Exd-Antp binding model (obtained by fitting a No Read Left Behind (NRLB) model (*23*) to the Lib-16 data set for Hth^HM^-Exd-Antp, ii) an Exd-Hth model (obtained by fitting a NRLB model to the Lib-16 data for Hth^FL^-Exd), and iii) a Hth-only model (PSAM model derived from Lib-30, using TTGACAGC as a seed). For each model view (in total there are [100-(number of positions specified by the model) +1] possible binding sites in each 100bp peak sequence), the score was computed for the “+” and strand respectively and only the maximum of the two was considered for each view. The cumulative peak score for each model was computed by summing up the scores across all views:

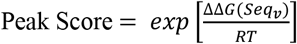

Testing for the stabilizing role of the Hth homeodomain to Exd-Hox sites was done by first considering the subset of peaks with a high confidence Exd-Hox site, a match to the consensus 12-mer NTGAYNNAYNNN (752 peaks). Next the subset of Exd-Hox peaks was split into the 30% of peaks with the strongest loss of Exd^−shape^ binding and the remaining 70% of less lost peaks. Both the cumulative Exd-Hox peak score as well as the affinity of the highest scoring Hth site (excluding the highest scoring Exd-Hox site) were compared between the two sets (t.test; 30% versus 70%). For the comparison between “high affinity” (Y5=T) and “low affinity” (Y5=C) sites, peaks were scanned for motif matches for NTGAY5NNAYNNN (752 peaks) and subdivided based on the identity of the Y5 position (T or C). The t-distribution was used to test for significant differences in the IP-coverage for Antp-GFP, Exd^WT^-V5, and Exd^−shape^-V5 between the two affinity classes.

### Clustering and peak to gene assignment

To cluster peaks based on their potential complex composition, 6 input features were considered: (i-iii) raw IP enrichment for Exd^WT^-V5, Exd^−shape^-V5, Antp^GFP^, and (iv-vi) peak scores for Exd-Antp, Exd-Hth (both NRLB models), and Hth-only (PSAM model). The resulting peak by feature table, was then transformed into standard scores (Z-scores) prior to cluster analysis. To cluster peaks on the 6 input features, the R package ‘flexclust’ was used with function cclust (method = “neuralgas”, k = 8). Eight clusters were chosen to allow for capturing of all possible complexes in addition to affinity differences and potentially unknown modes or accessibility driven, non-specific binding. Complex composition was assigned based on considering the average feature score for each cluster, as well as the degree of signal loss and peak accessibility (ATAC-seq; not included in the clustering).

### Hi-C data analysis

To visualize Hi-C contacts among Exd peak sets, only the 5Kb bins containing a peak were extracted, considering two options: i) in order to preserve one observation per peak, a 5kb bin for each peak was selected, resulting in duplicate Hi-C bins yet unique interaction counts per peak; ii) duplicate Hi-C bins were removed to test for the unique set of chromatin interactions spanned by Exd peak containing chromatin regions (Fig. S6A). In the same manner, random controls were generated by sub-sampling from the entire set of ATAC-seq peaks (~20,000). Random samples were size-matched and for the motifless peak set also accessibility-matched. The latter was achieved by matching the distribution of randomly chosen accessible sites to the true distribution of motifless peaks by restricting ATAC-seq sites to fall within the motifless [0.2,0.95] accessibility percentiles. To compute p-values, at least 50 such random size-matched (accessibility-matched) samples were generated. Average Hi-C contact frequencies were obtained by simply taking the mean across all 5Kb binned Hi-C contacts for a particular peak set. To generate an average Exd^−shape^ binding loss for motifless Exd binding sites based on their connectivity with classified Exd binding sites, three approaches were used: i) the Exd^−shape^ binding loss at the site with the highest interaction frequency with each specific motifless site was used; ii) the average Exd^−shape^ binding loss at one site per classified cluster (highest contact) was computed by using the log2 value of the interaction frequency with the motifless site as a weight; iii) the average Exd^−shape^ binding loss across all intra-chromosomal contacts between each motifless and all other classified Exd binding sites were computed by using the log2 value of the interaction frequency as a weight.

To assign peaks to a promoter the vanilla coverage (VC) normalized contact frequency for each peak across all promoters in the RNA-seq data set was computed. To simplify the analysis, only one promoter per gene (in case of multiple isoforms) was considered; the choice of promoter is based on whether a specific isoform was itself differentially expressed or (if not) whichever isoform had highest expression levels. The highest scoring peak-promoter interactions were then taken as the most likely target gene for each peak. To determine whether an individual promoter is significantly contacted by any of the five Exd peak types, the cumulative peak-promoter contact frequency within ± 50 Kb of the promoter was computed for each peak type separately. To determine which promoter had above expected contact frequency, p-values were computed based on a Wilcoxon test using the cumulative promoter-peak type contacts.

To test for a general connection between promoter-Exd peak interactions, the cumulative Exd peak-promoter VC contact frequency within ± 50 Kb of the promoter was extracted from the Hi-C data. Pearson correlation was next computed between the log2-gene-expression-fold change and the cumulative contact frequency. For visualization, promoters were split into equally sized bins (in 5 percent increments) based on the cumulative contact frequency.

### Gene Ontology analysis

Gene Ontology (GO) analysis was performed using the R package goseq (*24*). Tests were performed using the set of promoters that were both upregulated (5% FDR) and had an associated Exd peak, based on the maximum contact peak-to-promoter method described above. To test for contribution for individual complexes, only those promoters associated with a specific complex were used. Only GO categories with less than 1500 genes that scored significant in at least one of the complex-specific or all Exd gene sets (after correcting for multiple hypothesis testing) were considered for visualization. To test whether the enriched GO categories overlap with central nervous system (CNS)- or wing-specific functions, GO analysis was also performed on the intersection between upregulated genes in Exd^−shape^ and i) the geneset upregulated in the CNS or ii) in wing discs from a transcriptome comparison of wild-type tissues (for genotype information see above). If a GO category scored significant in one of the two latter genesets it was colored as CNS- or wing-specific respectively.

