## Supplemental Figures and Tables for "Context-dependent gene regulation by transcription factor complexes"

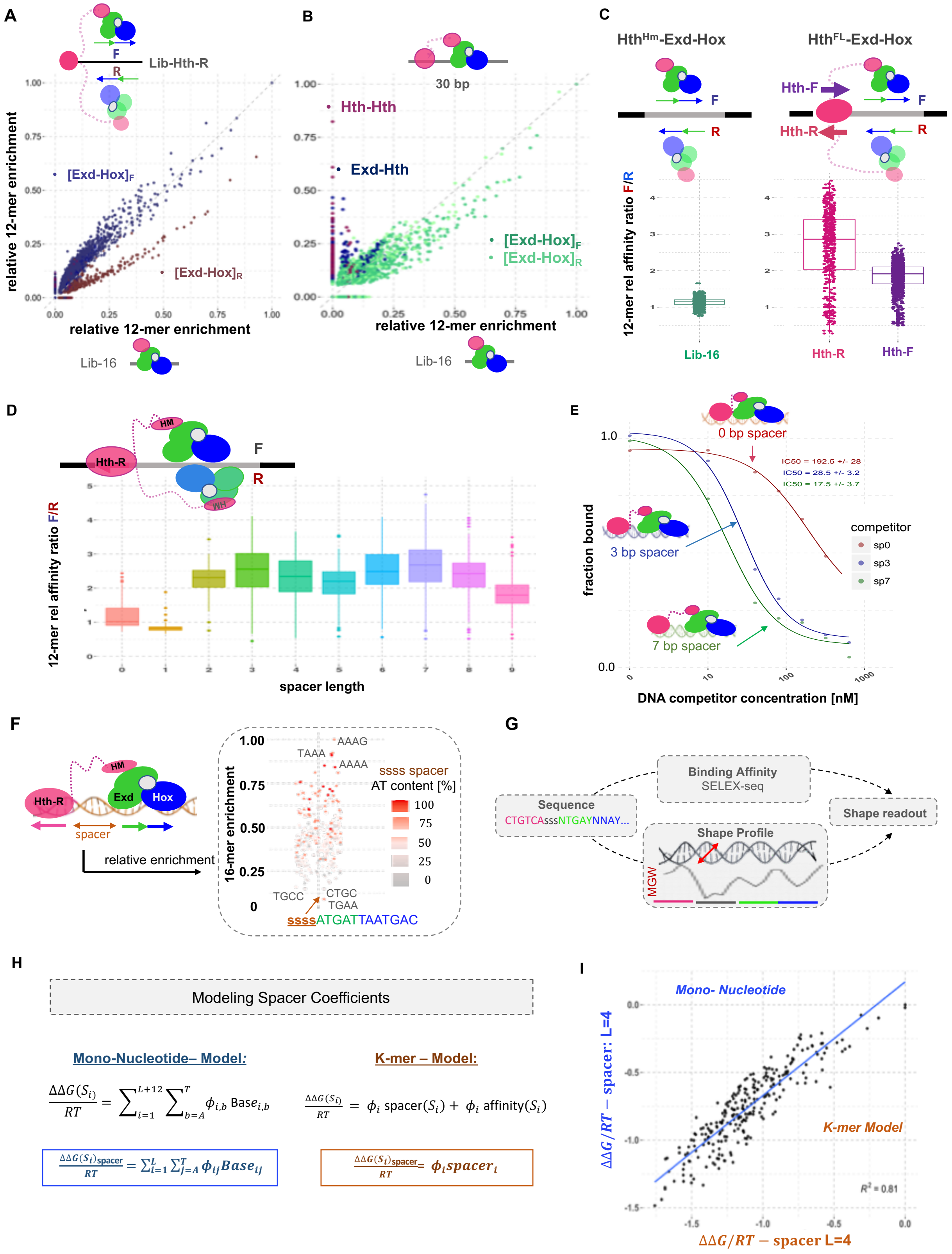
A
Lib-Hth-R
F
R
[Exd-Hox]F
[Exd-Hox]R
relative 12-mer enrichment
Lib-16
B
30 bp
Hth-Hth
Exd-Hth
[Exd-Hox]F
[Exd-Hox]R
relative 12-mer enrichment
Lib-16
C
HthHm-Exd-Hox
HthFL-Exd-Hox
Hth-F
Hth-R
12-mer rel affinity ratio F/R
Lib-16
Hth-R
Hth-F
D
HM
Hth-R
F
R
WH
12-mer rel affinity ratio F/R
spacer length
E
0 bp spacer
IC50 = 192.5 +/- 28
IC50 = 28.5 +/- 3.2
IC50 = 17.5 +/- 3.7
3 bp spacer
7 bp spacer
competitor
sp0
sp3
sp7
fraction bound
DNA competitor concentration [nM]
F
HM
Hth-R
Exd
Hox
spacer
relative enrichment
16-mer enrichment
AAAG
TAAA
AAAA
ssss spacer
AT content [%]
TGCC
CTGC
TGAA
ssssATGATTAATGAC
G
Binding Affinity
SELEX-seq
Sequence
CTGTCAsssNTGAYNNAY...
Shape Profile
MGW
Shape readout
H
Modeling Spacer Coefficients
Mono-Nucleotide– Model:
K-mer – Model:
I
Mono- Nucleotide
K-mer Model
ΔΔG/RT – spacer: L=4
ΔΔG/RT – spacer L=4

**Fig. S1. Binding Preferences and Model Comparison. (A)** Comparison of Exd-Hox binding preferences in the presence or absence of a Hth site. Normalized relative 12-mer enrichments for the ternary $Hth^{Fl}$-Exd-Hox (Lib-Hth-R) are compared to those for $Hth^{HM}$-Exd-Hox (Lib-16). 12-mers are colored based on the most likely Exd-Hox binding orientation with respect to the to the upstream fixed library flank: $[\text{Exd-Hox}]_F$ with a higher enrichment score for the forward 12-mer (blue); and $[\text{Exd-Hox}]_R$ with a higher score for the reverse orientation (red). The fixed Hth site introduces a preference for the $[\text{Exd-Hox}]_F$ orientation. **(B)** 12-mer relative enrichments are compared between Lib-30 with trimeric $Hth^{Fl}$-Exd-Dfd and Lib-16 with $Hth^{HM}$-Exd-Dfd bound. Three main binding modes are recognized: i) Hth-dimer sequences (purple), ii) canonical Exd-Hth sites (dark blue) and ii) Exd-Hox sites (green; shade implies the orientation of the binding site with respect to the library sequencing flanks). **(C)** Effect size of orientation preferences introduced by the fixed Hth site in either Lib-Hth-F or Lib-Hth-R compared to Lib-16 $Hth^{Hm}$-Exd-Dfd. Binding to the forward or reverse $[\text{Exd-Hox}]_{(F/R)}$ motif occurs in equal ratios in the symmetric Lib-16 ($Hth^{HM}$-Exd-Hox), whereas both Hth-F and Hth-R libraries have relative enrichment ratios of $[\text{Exd-Hox}]_F$ over $[\text{Exd-Hox}]_R$ greater than one (average of ~ 3 for Lib-Hth-R, pink; of ~2 for Lib-Hth-F, purple). **(D)** Relative 12-mer enrichments by offset to fixed Hth site in Lib-HthR. Preference for $[\text{Exd-Hox}]_F$ varies with spacer length. **(E)** Competition assay validating the spacer preference of the trimeric $Hth^{FL}$-Exd-Hox complex: three different spacer sequences – 0, 3, and 7bp – in between the Hth and Exd-Hox binding sites (CTGTCA-$(N)_L$-ATGATTAATGAC) were used to compete with a radio-labeled Hth-Exd-Hox probe (CTGTCA-AAA-ATGATTAATGAC). Agreeing with the model (**Fig. 2A**), spacers of 3 and 7 bp competed out the labeled probe at lower concentrations compared to a suboptimal spacer of 0 bp (IC50 values of $17.5 \pm 3.7; 28.5 \pm 3.2,$ and $192.5 \pm 28$ for spacers of 7, 3 and 0 bp respectively). **(F)** Sequence preferences of the DNA spacer between Hth and Exd-Hox. Relative enrichment for 16-mers with a fixed Exd-Hox site at positions 5-16 are shown and colored based on A/T content. More enriched sequences tend to have a higher A/T content. **(G)** Schematic for detecting shape-readout of DNA spacer sequence: For a specific spacer length, mononucleotide models are fit to the SELEX round 2 count data and the affinity score and the minor grove width (MGW) is computed for each sequence. Intrinsic, average DNA MGW profiles are correlated with TF binding selectivity to test for shape readout. **(H)** Spacer scores were generated by either fitting a mono-nucleotide model (blue) or a k-mer model (orange) to the round-2 SELEX-seq count data. Either sum of coefficients (mono-nucleotide model), or the individual spacer coefficients were used to compute spacer affinity scores. **(I)** The 4-bp spacer $\Delta\Delta G/\text{RT}$ coefficients were compared for either the mono-nucleotide model (blue) or the k-mer model (orange) ($R^2$= 0.81).

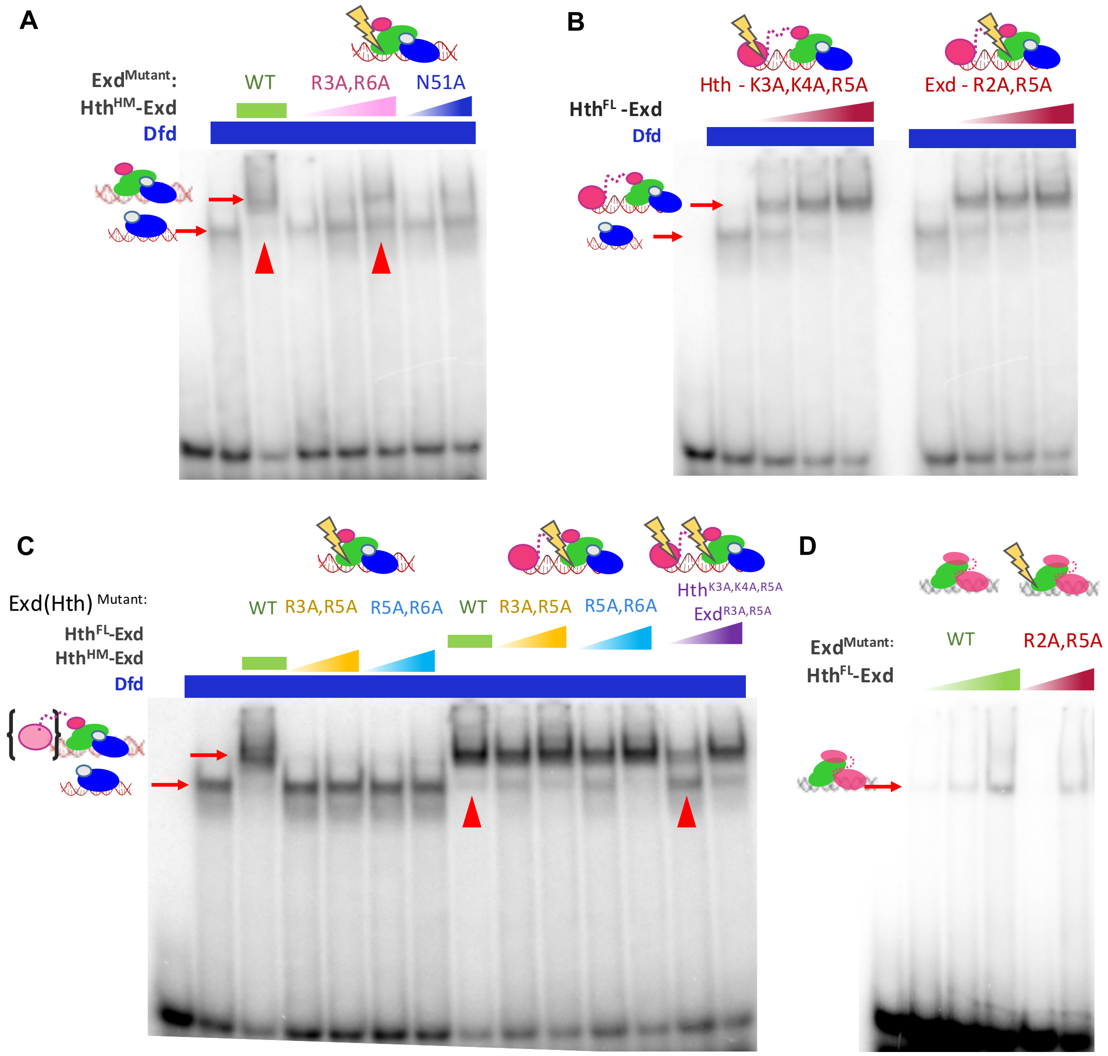


**Fig.S2. Binding assay for Hth and Exd mutant proteins. (A)** Electromobility shift assay for $Hth^{HM}$-Exd-Dfd with Exd double mutant protein (R3A,R6A; containing the two arginines that individually did not cause a loss of binding for the $Hth^{HM}$-Exd-Hox complex. Wild type Exd and the hydrogen-bond disrupting $Exd^{N51A}$ proteins are used for comparison: Lane 1: DNA only; lane 2: Dfd only, lane 3: $Hth^{HM}$-$Exd^{WT}$-Dfd, lane 4-6:$Hth^{HM}$-$Exd^{R3A,R6A}$-Dfd, lane 7-8: $Hth^{HM}$-$Exd^{N51A}$-Dfd. **(B)** Electromobility shift assay for $Hth^{-shape}$ and $Exd^{-shape}$ mutant proteins as used for the mutant Lib-Hth-(F/R) SELEX experiments. Lane 1: DNA only; lane 2&7: Dfd-only; lane 3-5: $Hth^{-shape}$-Exd-Dfd ($Hth^{K3A,K4A,R5A}$); lane 8-10: $Hth^{FL}$-$Exd^{-shape}$-Dfd ($Exd^{R2A, R5A}$). **(C)** Electromobility shift assay for combinations of Exd N-terminal arginine mutants with either $Hth^{HM}$ or $Hth^{Fl}$ isoform or with $Hth^{-shape}$ mutant protein. Lane 1: DNA only; lane 2: Dfd only, lane 3: $Hth^{HM}$-Exd-Dfd, lane 4-5: $Hth^{HM}$-$Exd^{(R3A,R5A)}$-Dfd, lane 6-7: $Hth^{HM}$-$Exd^{(R5A,R6A)}$-Dfd, lane 8: $Hth^{FL}$-Exd-Dfd; lane 9-10: $Hth^{FL}$-$Exd^{(R3A,R5A)}$-Dfd; lane 11-12: $Hth^{FL}$-$Exd^{(R5A, R6A)}$-Dfd, lane 13-14: $Hth^{K3A,K4A,R5A}$-$Exd^{R2A,R5A}$-Dfd. **(D)** Electromobility shift assay for $Exd^{R2A,R5A}$ in complex with $Hth^{FL}$. Lane 1: DNA only; lane 2-4: $Exd^{WT}$-$Hth^{FL}$ (concentration = 10, 50, 100 nM); lane 5-6: $Exd^{R2A,R5A}$-$Hth^{FL}$ (concentration = 10, 100 nM).

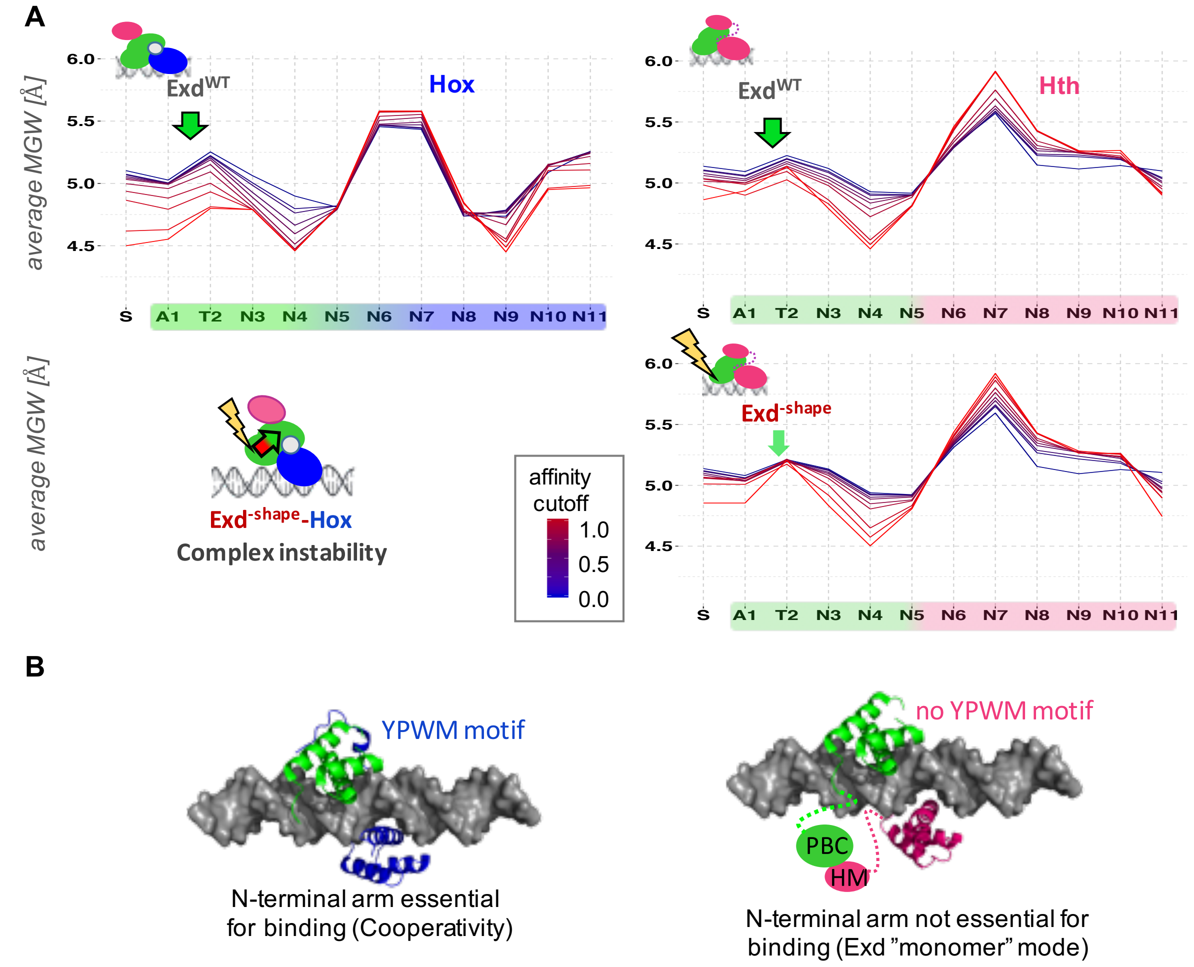


**Fig. S3. Exd's MGW readout is complex-dependent (A)** Comparison of wild-type and MGW-readout-mutant Exd (Exd$^{R2A,R5A}$ or Exd$^{-shape}$) in complex with either Hox (left panel) or Hth$^{FL}$ (right panel). The selection for a narrow DNA minor groove is attenuated in the wild-type Exd-Hth$^{FL}$ compared to Hth$^{HM}$-Exd-Hox. Mutating arginine 2 and 5 in Exd's N-terminal arm completely abrogates Exd-Hox complex binding, whereas it only has a minor impact on Exd-Hth$^{FL}$ binding further restricted to the first 2 positions (AT) along the Exd DNA interface (compare Figure 2 for Hth$^{FL}$-Exd-Hox). **(B)** Visualizing binding mode differences by structural superimposition of either Exd-Hox (PBD-ID: 2R5Y; Exd=green, Hox=blue) or Exd and MEIS1 (PDB-ID: 4XRM; pink) onto B-DNA. In Exd-Hox the YPWM motif directly interacts with Exd's HD, potentially intensifying Exd's shape readout through cooperative binding behavior. Hth uses its HM-domain that is connected via a flexible linker to Exd's PBC domain (indicated here as colored circles) to connect to Exd. It is likely that the binding is DNA mediated thus allowing Exd to bind in its native, monomeric mode.

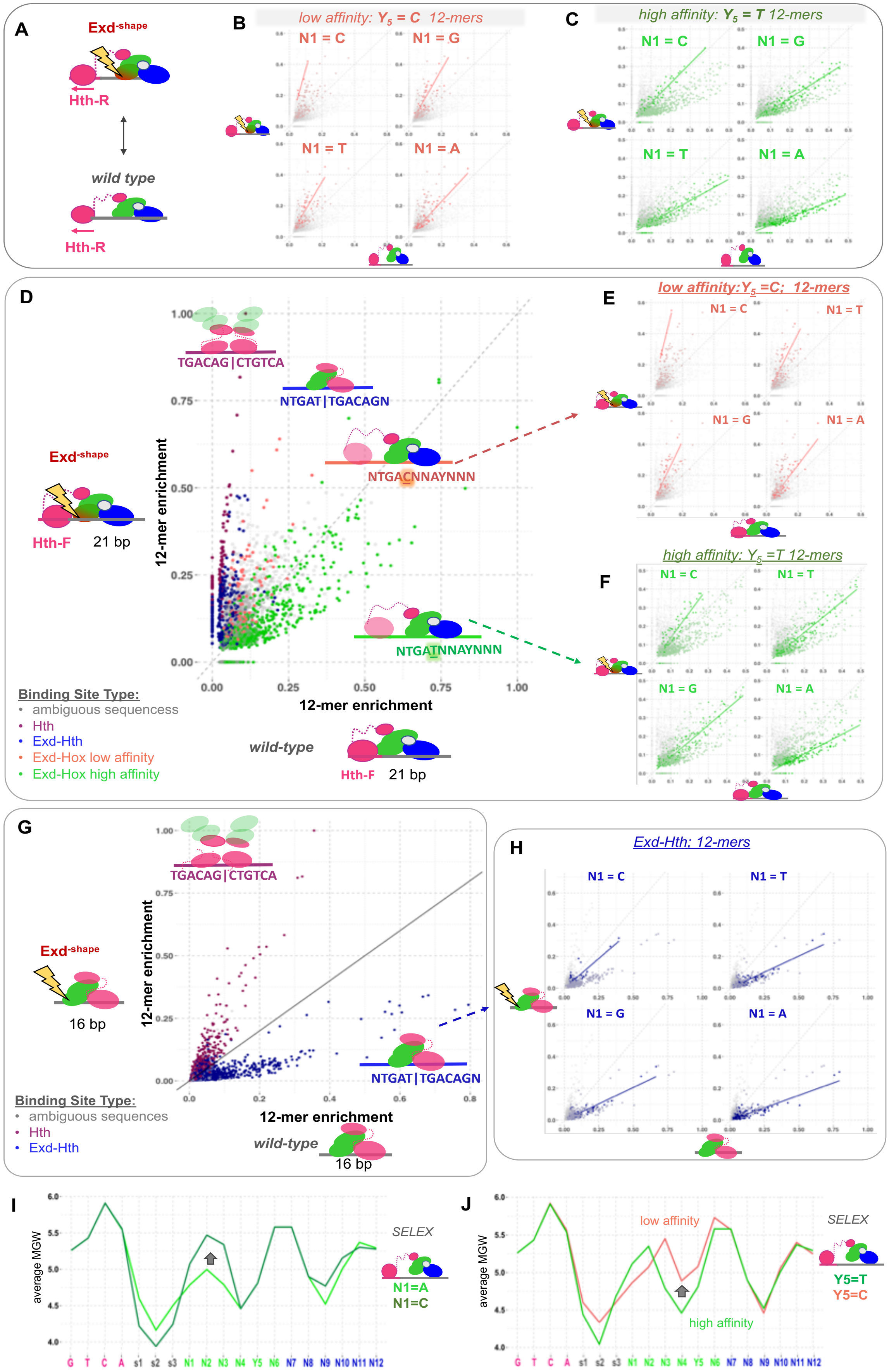
A
Exd-shape
Hth-R
wild type
Hth-R
B
low affinity: $Y_5$ = C 12-mers
N1 = C
N1 = G
N1 = T
N1 = A
C
high affinity: $Y_5$ = T 12-mers
N1 = C
N1 = G
N1 = T
N1 = A
D
TGACAG|CTGTCA
NTGAT|TGACAGN
NTGACNNAYNNN
NTGATNNAYNNN
Exd-shape
Hth-F
21 bp
12-mer enrichment
12-mer enrichment
Binding Site Type:
ambiguous sequencess
Hth
Exd-Hth
Exd-Hox low affinity
Exd-Hox high affinity
wild-type
Hth-F
21 bp
E
low affinity:$Y_5$ =C; 12-mers
N1 = C
N1 = T
N1 = G
N1 = A
F
high affinity: $Y_5$ =T 12-mers
N1 = C
N1 = T
N1 = G
N1 = A
G
TGACAG|CTGTCA
Exd-shape
16 bp
12-mer enrichment
12-mer enrichment
NTGAT|TGACAGN
Binding Site Type:
ambiguous sequences
Hth
Exd-Hth
wild-type
16 bp
H
Exd-Hth; 12-mers
N1 = C
N1 = T
N1 = G
N1 = A
I
average MGW
SELEX
N1=A
N1=C
G T C A s1 s2 s3 N1 N2 N3 N4 Y5 N6 N7 N8 N9 N10 N11 N12
J
average MGW
low affinity
high affinity
SELEX
Y5=T
Y5=C
G T C A s1 s2 s3 N1 N2 N3 N4 Y5 N6 N7 N8 N9 N10 N11 N12

**Fig. S4. Exd$^{–shape}$ differentiates complex compositions. (A)** Comparison of 12-mer relative sequence enrichments between Hth$^{FL}$-Exd-Hox and Hth$^{FL}$-Exd$^{–shape}$-Hox (Lib-Hth-F).**(B)** and **(C)** Blow-up of Y5=C (top) or Y5=T (bottom) 12-mer sequences from Fig. 3A. Each individual box shows subsequences that vary in the N1 base identity (N1=A,C,G, or T) and the degree to which they are lost in Exd$^{–shape}$ (lines represent linear fit). **(D)** Comparison of 12-mer relative sequence enrichments between Hth$^{FL}$-Exd-Hox and Hth$^{FL}$-Exd$^{–shape}$-Hox (Lib-Hth-F). 12-mers are colored by the PSAM sequence score most likely explaining their enrichment: Hth-dimers (purple), canonical Exd-Hth$^{FL}$ binding (dark blue), Hth$^{FL}$-Exd-Hox bound to the lower affinity sites (Y5=C: NTGA**C**NNAYNNN; coral red), and Hth$^{FL}$-Exd-Hox bound to the high affinity sites (Y5=T: NTGA**T**NNAYNNN; green). **(E)** and **(F)** Blow-up of Y5=C (top) or Y5=T (bottom) 12-mer sequences from plot A. Each individual box shows subsequences that vary in the N1 base identity (N1=A,C,G, or T) and the degree to which they are lost in Exd$^{-shape}$ (lines represent linear fit). **(G)** Comparison of 12-mer relative sequence enrichments between Hth$^{FL}$-Exd and Hth$^{FL}$-Exd$^{–shape}$ (Lib-16). 12-mers are colored by the PSAM sequence score most likely explaining their enrichment: Hth-dimers (purple), canonical Exd-Hth$^{FL}$ binding (dark blue). **(H)** Blow-up of 12-mer sequences from plot (G). Each individual box shows subsequences that vary in the N1 base identity (N1=A,C,G, or T) and the degree to which they are lost in Exd$^{-shape}$ (lines represent linear fit). **(I)** Proposed mechanisms for the sequence-dependent, differential binding loss of Hth$^{FL}$-Exd$^{–shape}$ –Hox. Average MGW profiles are shown for the top 10 highest scoring sequences when the N1 base is either fixed to A (light green) or C (dark green) before fitting the binding model. N1=A type sequences have an optimal, narrow MGW along the entire N-terminal end of Exd, thus more likely to be affected by a mutation removing shape readout, whereas N1=C type sequences have a much wider MG thus their binding mode is less impacted. **(J)** As in **(I)** but for position Y5 (coral red: Y5=C & green Y5=T). Widening of the MGW at positions N3-6 and the resulting shift in readout preferences is likely to explain the differences in Exd$^{–shape}$ binding to the two types of sequences.

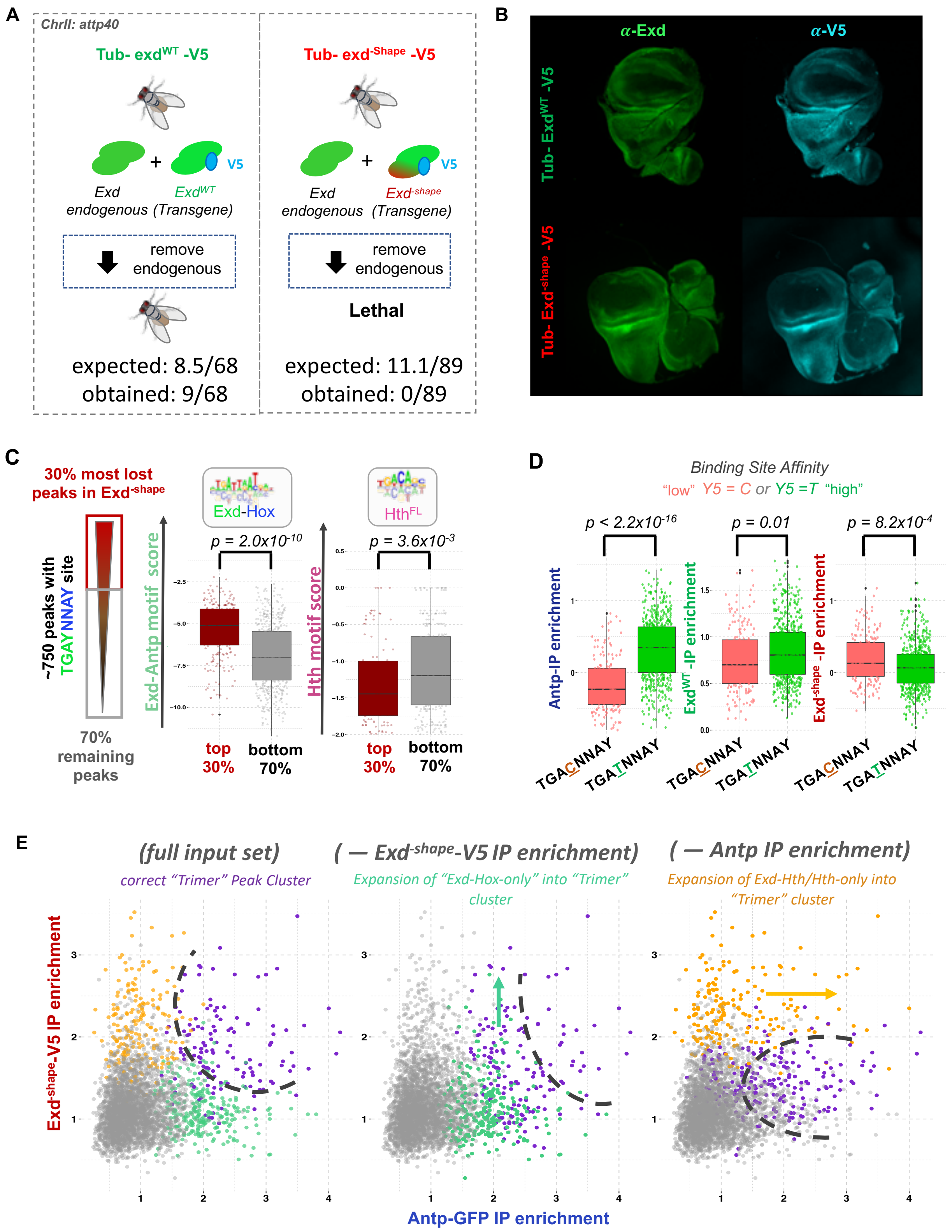
A
ChrII: attp40
Tub- exdWT -V5
Exd endogenous
ExdWT (Transgene)
V5
remove endogenous
expected: 8.5/68
obtained: 9/68
Tub- exd-Shape -V5
Exd-shape (Transgene)
Lethal
expected: 11.1/89
obtained: 0/89
B
α-Exd
α-V5
Tub- ExdWT -V5
Tub- Exd-shape -V5
C
30% most lost peaks in Exd-shape
~750 peaks with TGAYNNAY site
70% remaining peaks
Exd-Hox
HthFL
Exd-Antp motif score
Hth motif score
p = 2.0x10-10
p = 3.6x10-3
top 30%
bottom 70%
D
Binding Site Affinity
"low" Y5 = C or Y5 =T "high"
p < 2.2x10-16
p = 0.01
p = 8.2x10-4
Antp-IP enrichment
ExdWT -IP enrichment
Exd-shape -IP enrichment
TGACNNAY
TGATNNAY
E
(full input set)
correct "Trimer" Peak Cluster
( — Exd-shape-V5 IP enrichment)
Expansion of "Exd-Hox-only" into "Trimer" cluster
( — Antp IP enrichment)
Expansion of Exd-Hth/Hth-only into "Trimer" cluster
Exd-shape-V5 IP enrichment
Antp-GFP IP enrichment

**Fig. S5. Using Exd$^{-shape}$ to probe complex composition and binding mechanisms in vivo.**
**(A)** Transgenes carrying either Exd$^{Wt}$ or Exd$^{-shape}$ tagged with V5 are inserted into the attp40 landing site on chromosome II. Removing endogenous Exd transcripts is rescued by Exd$^{Wt}$-V5 (9 males of 68 progeny, expected 8.5) but not Exd$^{-shape}$-V5 (0 males of 89 progeny, expected 11, $p$ = 0.004). **(B)** Anti-Exd (green) and anti-V5 (cyan) stain of homozygous Exd$^{Wt}$ or Exd$^{-shape}$ V5-tagged 3$^{rd}$ instar imaginal wing discs (in the background of endogenous Exd). V5 signal follows total Exd stain and is nuclear in both genotypes. **(C)** Hth$^{FL}$ binding stabilizes binding loss of Exd-Hox dimers: The 752 peaks containing a match to the consensus Exd-Hox site (TGAYNNAY) were split according to Exd$^{-shape}$ binding loss (ratio Exd$^{WT}$/Exd$^{-shape}$; 30:70 - red:grey) and the peak sequence (100 bp around summit) was scored with either Exd-Antp (turquois) or Hth (pink) motif models. 30:70 peak sets were tested for differences in Exd-Antp BS peak score and the presence of a secondary Hth motif. **(D)** The 752 peaks with a match to an Exd-Hox TGAYNNAY site were split into two groups $Y_5$=C (low affinity; coral red) and $Y_5$=T (high affinity; green), and the *in vivo* sequence selectivity for Antp, Exd$^{WT}$ and Exd$^{-shape}$ is compared between the two groups. Exd$^{-shape}$ prefers the low affinity $Y_5$=C over the high affinity $Y_5$=T sites. **(E)** Scatterplot of Exd$^{-shape}$-V5 IP enrichment against Hox (Antp-GFP) IP enrichment for all ~3700 Exd$^{WT-}$V5 peak summits. Coloring is based on peak clustering using i) the full feature set (three IP signals (Exd$^{WT}$-V5, Exd$^{-shape}$-V5, Antp-GFP) and three binding site scores (Exd-Antp, Hth-only, Exd-Hth; *in vitro* binding models); ii) 5 out of 6 features, leaving out Exd$^{-shape}$-V5 (middle panel); iii) 5 out of 6 features, leaving out Antp IP signal (right panel). Only the full model correctly differentiates the "trimeric" cluster that shows stabilized Exd$^{-shape}$ signal at high Antp IP signal (purple). Both reduced feature sets produce impure clustering; with leaving out Exd$^{-shape}$-V5 resulting in blending of "Exd-Hox-only" and "Trimer" sites and leaving out Antp in blending of "Exd-Hth/Hth-only" and "Trimer" sites (see dashed lines and arrows).

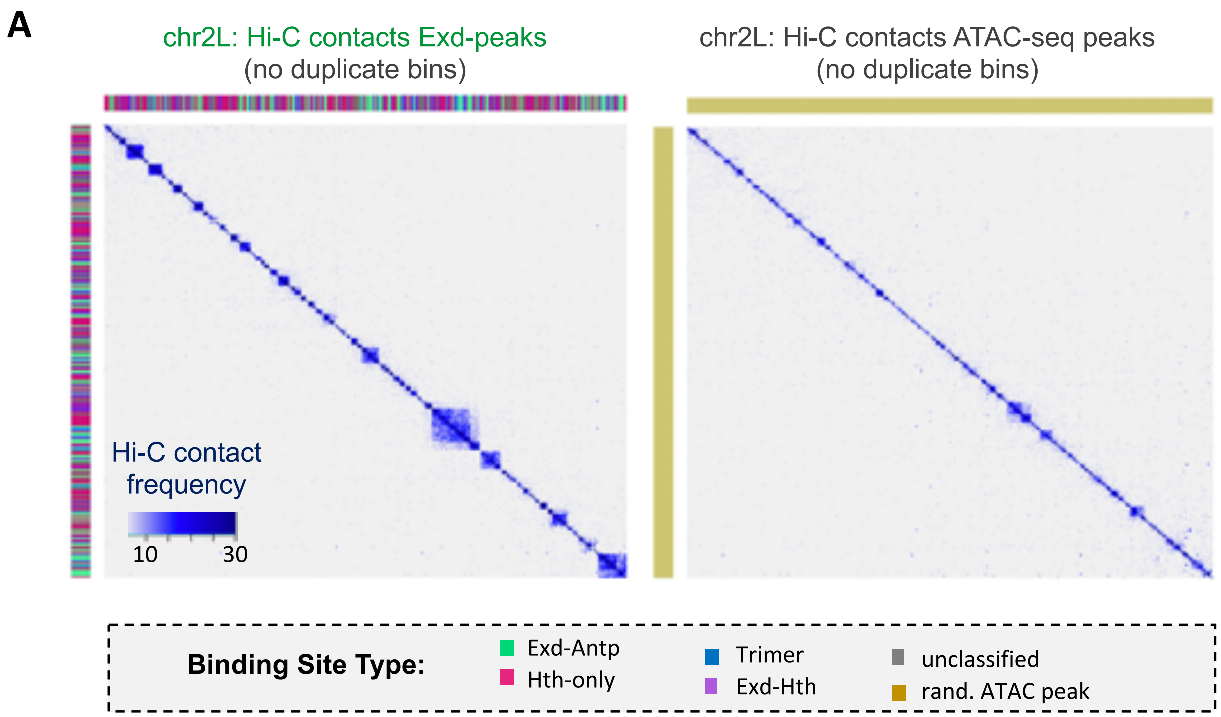


**Fig. S6. Exd binding sites are physically interacting. (A)** Hi-C contact maps of wild-type (incl. tub>exd$^{WT}$-V5 transgene) third instar larval wing discs isolating either the set of Exd peaks (left) or a size-matched random set of ATAC-seq peaks (right). Clusters of sites corresponding to distinct genomic contact domain structures are seen for the Exd peakset, but not for a size-matched random sample of accessible genomic regions (p.value = $2.6*10^{-32}$). Only unique Hi-C bins are retained; note that the choice of peak color in the left panel (beside and above the Hi-C map) occurs at random for all Hi-C bins that contain two or more Exd peaks.

A

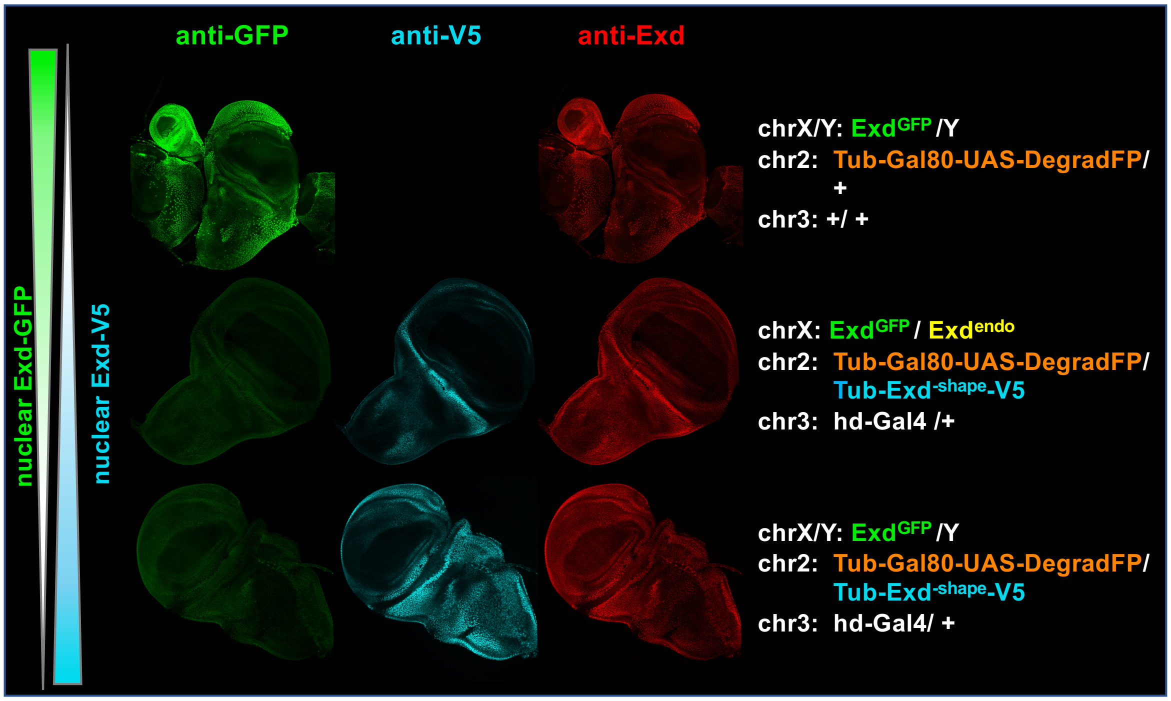
anti-GFP
anti-V5
anti-Exd
nuclear Exd-GFP
nuclear Exd-V5
chrX/Y: ExdGFP /Y
chr2: Tub-Gal80-UAS-DegradFP/ +
chr3: +/ +
chrX: ExdGFP / Exdendo
chr2: Tub-Gal80-UAS-DegradFP/ Tub-Exd-shape-V5
chr3: hd-Gal4 /+
chrX/Y: ExdGFP /Y
chr2: Tub-Gal80-UAS-DegradFP/ Tub-Exd-shape-V5
chr3: hd-Gal4/ +

B

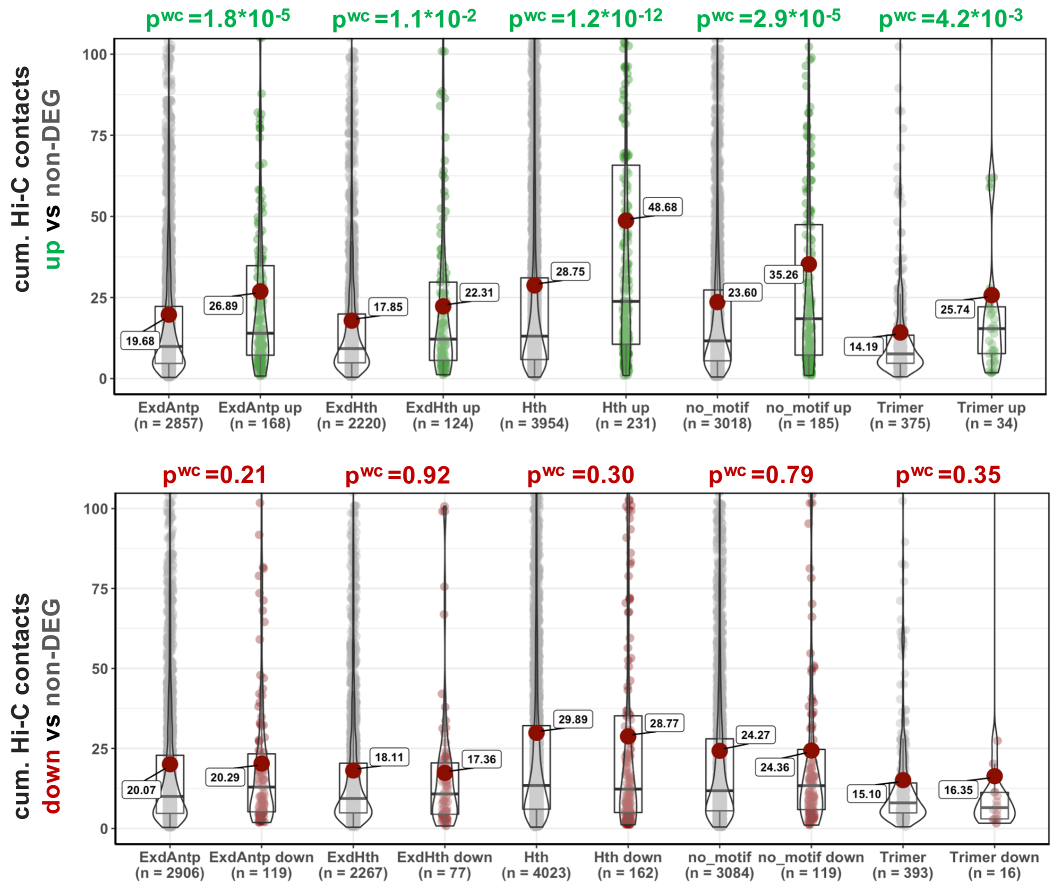
pwc =1.8*10-5
pwc =1.1*10-2
pwc =1.2*10-12
pwc =2.9*10-5
pwc =4.2*10-3
cum. Hi-C contacts up vs non-DEG
19.68
26.89
17.85
22.31
28.75
48.68
23.60
35.26
14.19
25.74
ExdAntp (n = 2857)
ExdAntp up (n = 168)
ExdHth (n = 2220)
ExdHth up (n = 124)
Hth (n = 3954)
Hth up (n = 231)
no_motif (n = 3018)
no_motif up (n = 185)
Trimer (n = 375)
Trimer up (n = 34)
pwc =0.21
pwc =0.92
pwc =0.30
pwc =0.79
pwc =0.35
cum. Hi-C contacts down vs non-DEG
20.07
20.29
18.11
17.36
29.89
28.77
24.27
24.36
15.10
16.35
ExdAntp (n = 2906)
ExdAntp down (n = 119)
ExdHth (n = 2267)
ExdHth down (n = 77)
Hth (n = 4023)
Hth down (n = 162)
no_motif (n = 3084)
no_motif down (n = 119)
Trimer (n = 393)
Trimer down (n = 16)

**Fig. S7. Exd$^{-shape}$ as a tool to study complex-specific function in vivo. (A)** Testing the deGradFP system: Images show a confocal plane of third instar imaginal wing discs, with larva raised at 18℃ and shifted to 29℃ 24h prior to dissection. Three different genotypes are shown: male wing disc containing one copy of Exd$^{GFP}$ on the X chromosome and the deGradFP System on the second chromosome (top); female wing disc containing one Exd$^{GFP}$ and one copy of endogenous Exd on the X chromosomes, the DegradFP system and the transgenic Exd$^{-shape}$-V5 on the 2$^{nd}$ chromosome, as well as the headcase-Gal4 (hd-Gal4) on the third chromosmes (middle); as middle, except male disc with the endogenous Exd copy absent (bottom). In the absence of hd-Gal4, the deGradFP system is not active and Exd$^{GFP}$ is nuclear (top). Upon activation of the deGradFP system the nuclear Exd$^{GFP}$ is greatly reduced and Exd$^{-shape}$-V5 competes with endogenous Exd for nuclear localization (middle). Upon removal of endogenous Exd (by only considering male flies), and continuing to remove Exd$^{GFP}$, transgenic Exd$^{-shape}$-V5 is the only source of Exd protein and nuclear levels increase. **(B)** Promoters of up, but not downregulated genes in Exd$^{-shape}$ vs Exd$^{WT}$ display a significantly higher Hi-C contact frequency (Wilcox test (wc)) for each of the five major peak classifications – Exd-Hox, Exd-Hth, Hth-only, motifless ("no-motif"), and Trimer binding sites – compared to promoters whose transcripts are not differentially expressed (non-DEG; FDR threshold set to 5%).

Table S1. **List of individual mutations made for Exd and Hth**

| Protein | Mut 1 | Mut 2 | Mut 3 | Mut 4 | Mut 5 | Mut 6 | Mut 7 | Mut 8 | Mut 9 |
|---|---|---|---|---|---|---|---|---|---|
| Exd | R2A | R3A | R5A | R6A | N51A | R2A & R5A | R3A & R5A | R5A & R6A | R3A & R6A |
| Hth | K4A & R5A | K3A & K4A & R5A | | | | | | | |

**Table S2. Full description of library sequences**

| | | | |
|---|---|---|---|
| Lib-16 design 1 | 5’ | GTTCAGAGTTCTACAGTCCGACGATCTGG **(16xN)** CCAGCTGTCGTATGCCGTCTTCTGCTTG | 3’ |
| Lib-16; design 2 | 5’ | GGTAGTGGAGGTGGGCCTGG **(16xN)** CCAGG GAGGTGGAGTAGG | 3’ |
| Lib-21 Hth-F | 5’ | GTTCAGAGTTCTACAGTCCGACGATC <u>**TGACAG**</u> **(21xN)** CCCGGGTCGTATGCCGTCTTCTGCTTG | 3’ |
| Lib-21 Hth-R | 5’ | GTTCAGAGTTCTACAGTCCGACGATC <u>**CTGTCA**</u> **(21xN)** CCCGGGTCGTATGCCGTCTTCTGCTTG | 3’ |
| Lib-30 | 5’ | GTTCAGAGTTCTACAGTCCGACGATCTGG **(30xN)** CCCGGGTCGTATGCCGTCTTCTGCTTG | 3’ |

Fixed Hth binding sites are underlined and in bold. Random region length in bold and in parentheses.
